# Real-Time Wide-Field Fluorescence Lifetime Imaging via Single-Snapshot Acquisition for Biomedical Applications

**DOI:** 10.1101/2025.04.16.646646

**Authors:** Vikas Pandey, Euan Millar, Ismail Erbas, Luis Chavez, Jack Radford, Isaiah Crosbourne, Mansa Madhusudan, Gregor G. Taylor, Nanxue Yuan, Claudio Bruschini, Stefan T. Radev, Margarida Barroso, Andrew Tobin, Xavier Michalet, Edoardo Charbon, Daniele Faccio, Xavier Intes

## Abstract

Fluorescence lifetime imaging (FLI) is a powerful tool for investigating molecular processes, microenvironmental parameters, and molecular interactions across tissue to (sub-)cellular levels. Despite its established value in numerous biomedical applications, conventional FLI techniques are hindered by long acquisition times. This limitation restricts their use in real-time scenarios, such as monitoring fast biological processes, studying live organisms, and in environments that require rapid imaging and immediate inference, such as clinical image-guided interventions. Here, we present a novel FLI approach that combines a large-format time-gated SPAD array with dual-gate acquisition capability, alongside a rapid lifetime determination algorithm. This integration allows for real-time fluorescence lifetime estimation through single-snapshot acquisitions, eliminating the need for traditional, time-consuming time-resolved data collection. We demonstrate the scalability and versatility of this method by achieving real-time FLI across challenging biomedical applications, ranging from capturing fast neural dynamics at the microscopic scale, performing multimodal 3D volumetric FLI of tumor organoids at the mesoscopic scale, to macroscale FLI in both direct and highly scattering regimes. Furthermore, we validate its utility in fluorescence lifetime-guided surgical procedures using tissue-mimicking phantoms. Overall, this new methodology significantly enhances the temporal and spatial capabilities of FLI, opening the door to the assessment of fast dynamic biomedical signals. It also enables the seamless integration of FLI into clinical workflows, particularly in applications like fluorescence-guided surgery.

## I. Introduction

Fluorescence lifetime imaging (FLI) is a powerful optical imaging technique that enables the investigation of cellular and molecular processes across various imaging scales – including metabolism, pH levels, cellular respiration, protein interactions, and drug-target engagement in living organisms and [1]. Its unique ability to provide contrast based on fluorescence decay dynamics makes it particularly valuable in translational applications, where it can assess pathological microenvironments and quantify/guide therapeutic interventions *in vivo*. However, FLI is inherently a computational imaging technique that relies on prolonged time-resolved data acquisition, followed by computationally expensive processing pipelines. Traditional methods, such as nonlinear least squares fitting (NLSF) and maximum likelihood estimation (MLE), are time-consuming and require significant computational resources, creating a substantial bottleneck in fast fluorescence lifetime image generation [2]. This limitation is particularly critical in applications where rapid inference is necessary, such as tracking fast biological processes or contributing to the development of methods for clinical feedback at the bedside, such as delineation of tumor margins during resections [3]. Consequently, FLI has remained largely confined to *in vitro* microscopic biological studies or preclinical research with limited adoption in clinical workflows.

Recent advances in computational power and deep learning techniques have significantly reduced the processing time required to estimate fluorescence lifetime decay contrast, reducing it from hours to (near) real time [4]–[10]. Despite these breakthroughs, the bottleneck in time-resolved data acquisition remains, particularly for live biological studies involving dynamic processes occurring on millisecond timescales. Capturing time-resolved data for such rapid events across large field-of-views and with high spatial resolution remains a significant challenge [11]. Achieving such performance requires advancements in both data acquisition technologies and computational approaches. To address this, we introduce a rapid lifetime determination (RLD) method based on a single-snapshot acquisition, realized by simultaneously acquiring time-gated and full-temporal aperture measurements.

In this study, we employ a single-camera solution featuring a novel dual-gated SPAD architecture, SwissSPAD3 (SS3) [12], which enables the simultaneous capture of both gated and full-intensity measurements in a single-snapshot. This capability ensures that both types of measurements are perfectly correlated, regardless of intensity variations caused by external factors such as illumination fluctuations, photobleaching, optical properties changes (like bleeding), or rapid biological events (breathing, pulsating flow, etc). When combined with the RLD algorithm, this system facilitates accurate fluorescence lifetime estimation using a single-snapshot acquisition. As a result, it eliminates the need for traditional time-resolved data collection, streamlining the imaging process while maintaining high precision. We demonstrate the versatility of our approach by achieving near-real-time FLI at rates of at least 5 frames per second (fps) across a wide range of imaging scales. These include monitoring fast neuronal signals at the microscopic scale through microscopic FLI, performing multimodal 3D volumetric FLI of a large tumor organoid using a mesoscopic lightsheet illumination setup (mesoscopic FLI), and performing large-area FLI (8 cm × 6 cm) for near-infrared fluorophores in both direct and highly scattering optical imaging regimes (macroscopic FLI).

Furthermore, we validate the efficacy of our real-time FLI method in a mock fluorescence lifetime-guided surgical procedure using tissue-mimicking phantoms modeled after mouse and human breast anatomy. Our findings establish a strong foundation for integrating real-time FLI into diverse micro-, meso-, and macroscopic applications. Notably, we demonstrate its potential for clinical integration, with significant promise for applications such as the precise identification of tumor margins in image-guided surgery [13], [14], thus facilitating broader clinical adoption of FLI.

## II. Results

### A. Real-Time Macroscopic FLI: Direct and Diffuse Optical Imaging

To demonstrate the implementation of real-time FLI in a macroscopic imaging setup with a large field-of-view (8 cm × 6 cm), we used the near-infrared (NIR) dye Alexa Fluor 700 (AF700) dissolved in two different solvents: phosphatebuffered saline (PBS) and dimethyl sulfoxide (DMSO). The NIR-I dye and solvents were selected primarily for two reasons: to produce distinct fluorescence lifetimes due to the different solvent polarities, and because we have previously demonstrated the use of these NIR-I dyes in a non-invasive drug-target quantification study [15], [16]. 1 *µ*M solutions of AF700 in PBS and DMSO were prepared and placed in two separate wells of a 96-well plate.

Dual-gate single-snapshot data were captured using SS3 in continuous acquisition mode, in which a series of 255 or 1020 pairs of one-bit images are accumulated on board and transferred to the computer sequentially. Each pair consists of an intensity image (INT), representing the number of detected photons (0 or 1) during the 1-bit integration period (*>* 20.4 *µs*), and a gated image (G2), capturing the number of photons (0 or 1) recorded within the user-defined “gate” within each laser period. Figure 1(a) shows the schematic of our macroscopic FLI setup (see subsubsection IV-E.3) used to capture data, while Figure 1(b) illustrates the pulsed illumination, first-order decay of sample fluorescence, and the SS3 dual-gate signal acquisition mechanism. The RLD algorithm, described in subsection IV-C, was implemented in the acquisition software (see Supplementary Section 2 for details) to generate FLI visualization in real time. The dual-gate acquisition principle and additional details of SS3 can be found in subsection IV-A. For this experiment, the G2 images were recorded with a gate width (*W*) of 3 ns, and the gate delay was selected at 5.5 ns. The details of gate selection are explained in **Supplementary Section 4.3**.

**Fig. 1:**
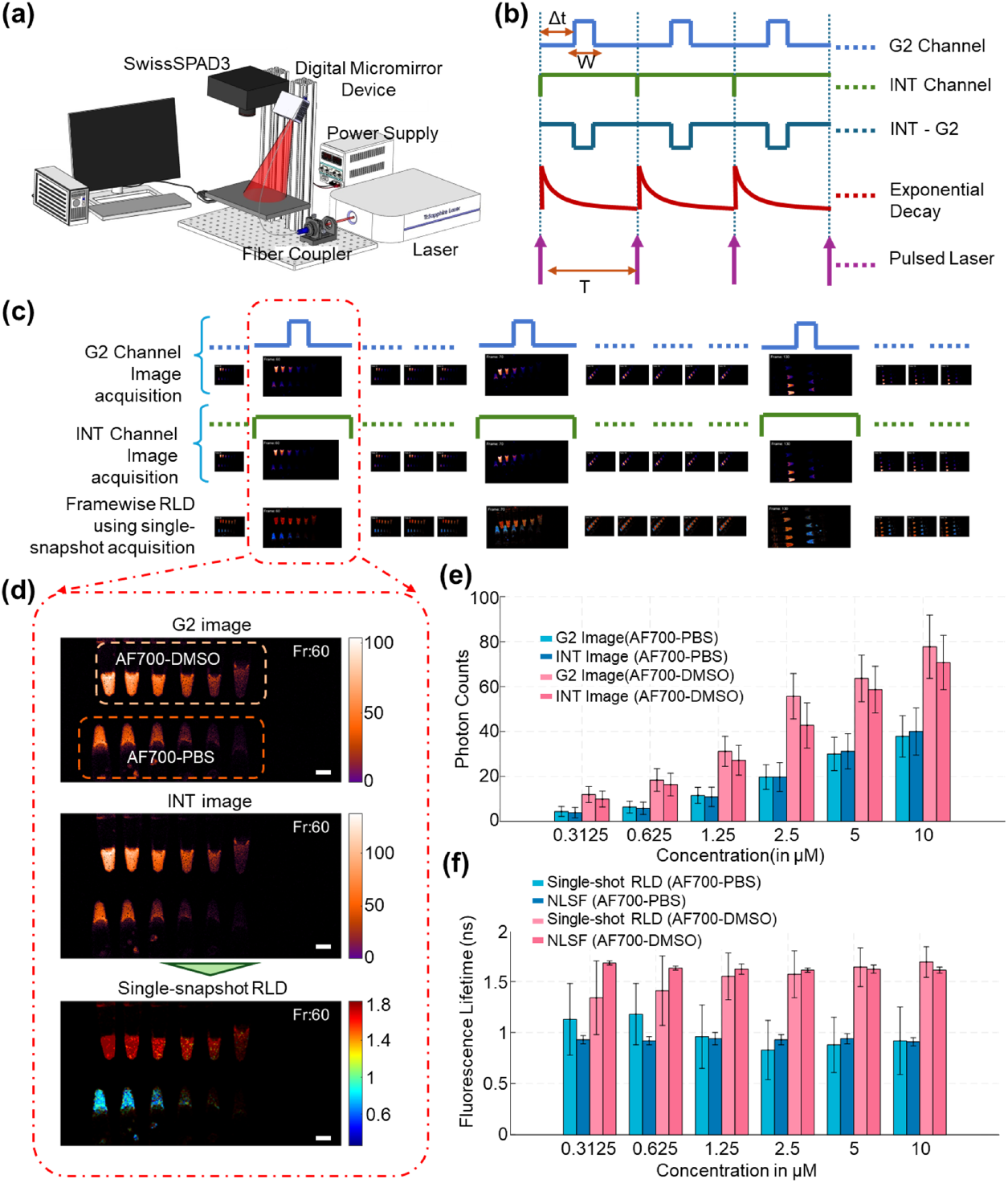
Real-time Macroscopic Fluorescence Lifetime Imaging: (a) Schematic of the macroscopic FLI setup, comprising an 80 MHz tunable pulsed laser, a digital micromirror device (DMD) for structured or wide-field illumination, and a time-resolved SwissSPAD3 (SS3) camera. (b) Illustration of SS3 dual-gate signal acquisition: the fluorescence sample is excited using a pulsed laser at a repetition rate of 80 MHz, and the sample fluorescence decay (SFD) follows first-order kinetics. The G2 channel records SFD signals at a user-selected gate width (W) and delay (Δt) relative to the illumination pulse, while the INT channel records the SFD signal over the entire laser period T. The INT-G2 signal can be derived for computational analysis. (c) Schematic illustrating a sequence of time frames showing G2 and INT images acquisition first and second row, respectively, with their respective gating mechanisms on top of the frame and the corresponding real-time lifetime computations in the third row. Three randomly selected frames are shown enlarged to emphasize sample movement and real-time FLI performance. Refer to Supplementary Video 1 for the full dynamic visualization. (d) Enlarged view of frame 60, showing detailed G2 and INT images alongside the computed fluorescence lifetime map. The sample contains Alexa Fluor 700 (AF700) dye in two sets of six micro-scale plastic tubes, each set dissolved in either PBS or DMSO. The dye concentrations range from 10 *µ*M to 0.3125 *µ*M (10, 5, 2.5, 1.25, 0.625, and 0.3125 *µ*M) from left to right. The white scale bar represents 10 mm. (e) Variation in photon counts in the G2 and INT images as a function of AF700 dye concentration. (f) Comparison of single-shot RLD-estimated fluorescence lifetime with nonlinear least squares fitting (NLSF) across varying concentrations of AF700 in PBS and DMSO.

To demonstrate the real-time FLI computation capability, varying concentrations of AF700 (10 *µ*M, 5 *µ*M, 2.5 *µ*M, 1.25 *µ*M, 0.625 *µ*M, and 0.3125 *µ*M) dissolved in PBS and DMSO, were filled into two sets of six connected micro-scale plastic tubes (height 32 mm, volume capacity 0.5 mL).

AF700 exhibits significantly different fluorescence lifetimes of 1 ns and 1.65 ns in PBS and DMSO, respectively, as estimated using NLSF. Both sets of tubes were moved randomly within the macroscopic imaging field-of-view while acquiring continuous single-snapshot measurements and generating their corresponding real-time lifetime images. The different positions of each set of tubes in randomly selected frames are shown in Figure 1(c). Three randomly selected intermediate time points of the time-series in Figure 1(c) are shown at a larger scale in the strip to emphasize the movement of the sample during acquisition. The bottom strip of Figure 1(c) presents the corresponding single-snapshot RLD-based lifetime maps computed in real-time. The time-series signal capture and the simultaneous real-time lifetime computation of this experiment are shared in the video **Supplementary Video 1**.

Both sets of tubes exhibited a large range of photon counts (3 - 150), with higher photon counts in DMSO than PBS at the same concentration. However, solely using intensity information, these mixed tubes cannot be distinguished without prior knowledge. In contrast, their corresponding lifetime maps clearly differentiate the two solvent environments. The varying concentration of the AF700 affects the photon counts (Figure 1(e)), but the fluorescence lifetime remains unchanged (Figure 1(f)). At very low photon counts (*<* 10 photons per pixel per acquisition), specifically for AF700-PBS at the concentrations of 1.25 *µ*M, 0.625 *µ*M, and 0.3125 *µ*M, and for AF700-DMSO at 0.625 *µ*M and 0.3125 *µ*M, the tubes are only partially visible in G2 and INT images. At such low photon counts, it becomes challenging to distinguish the signals from noise and hence leads to over- or underestimation of lifetime. In this low-photon regime, single-snapshot RLD overestimated lifetimes by ∼20% in AF700-PBS and underestimated them by ∼15% in AF700-DMSO compared to NLSF. It is important to emphasize that NLSF methods use the summation of multiple full decay acquisitions of the sample, requiring prolonged acquisition times as well as pre-processing prior to NLSF implementation. At very low photon counts, single-snapshot RLD also requires further validation with other methods or enhancement using deep learning-based approaches.

The video of the aforementioned experiment validates realtime FLI through single-snapshot acquisition. The single-snapshot lifetime estimation was compared with the traditional NLSF method (see Supplementary Section 3) using full time-resolved acquisition. The lifetimes computed using NLSF were found to be in good agreement (Supplementary Figure 4(h)), confirming the accuracy of the single-snapshot RLD method.

The single-snapshot RLD method was then evaluated for a macroscopic FLI experiment in preclinical settings. Diffuse FLI is increasingly employed in non-invasive *in vivo* small animal imaging to validate new fluorescence probes designed for clinical use or to assess new targeted drug efficacy. Notably, our group pioneered the use of FLI for the quantification of probe-target/drug-tumor interactions *in vivo*, a critical parame-ter of drug action which could be assessed only using invasive methods [15].

A mouse-shaped tissue-mimicking phantom was built for this experiment, as detailed in subsubsection IV-D.2. Two Eppendorf tubes (0.5 mL) each containing 10 *µ*M AF700 dissolved in PBS and DMSO, respectively, were embedded at a depth of 2–3 mm from the phantom’s upper surface (in supine position). The mouse-shaped phantom was imaged in supine orientation (Figure 2(a)).

**Fig. 2:**
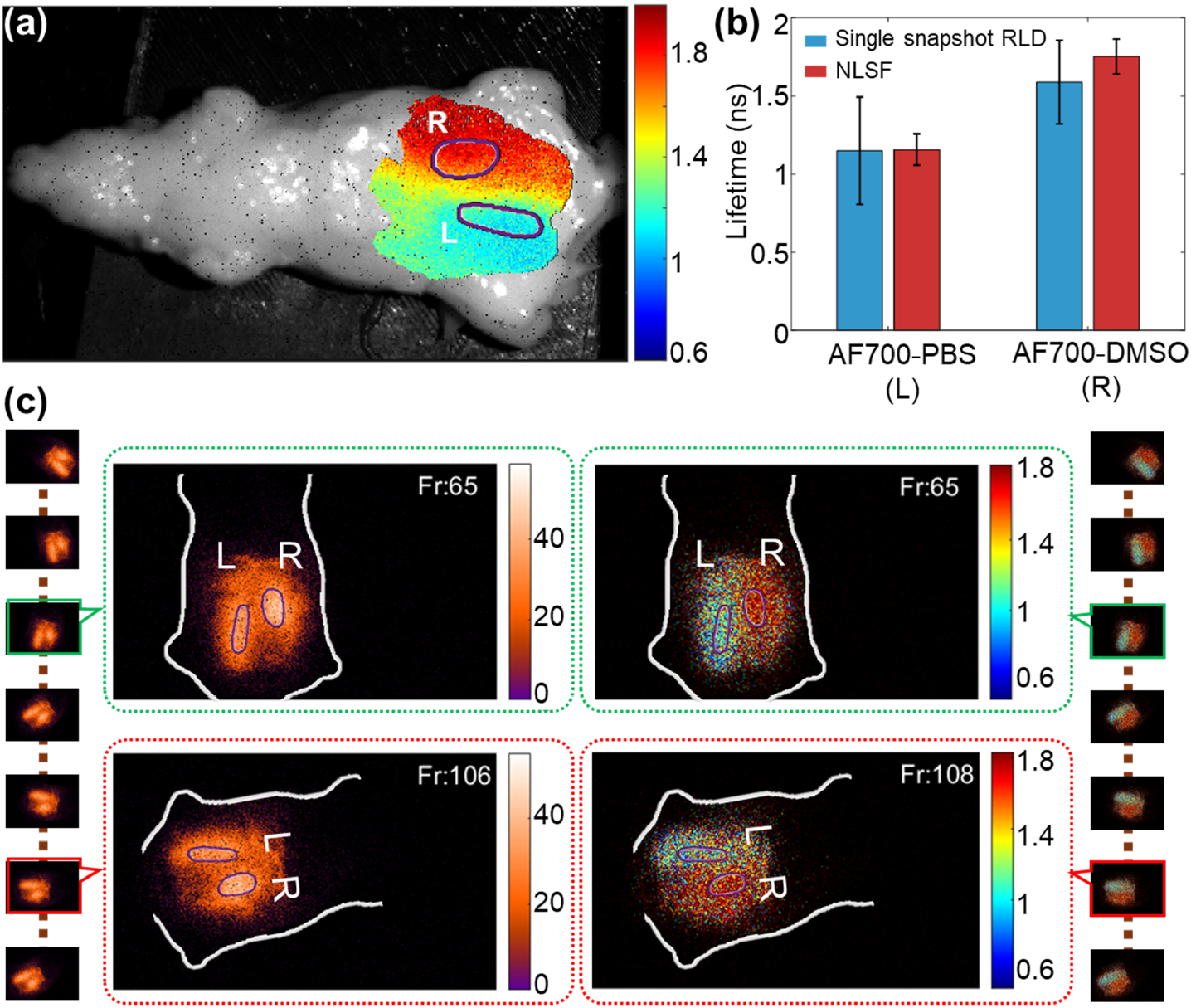
Macroscopic Diffuse Optical Fluorescence Lifetime Imaging. A tissue-mimicking, mouse-shaped phantom was imaged in a wide-field macroscopic FLI set-up (field-of-view: 8 cm × 6 cm). Two Eppendorf tubes (height: 32 mm, volume capacity: 0.5 mL) containing 10 *µ*M Alexa Fluor 700 dye in PBS (L) and DMSO (R) were embedded at a depth of 2–3 mm from the top surface. The model was positioned in a supine orientation for imaging. (a) NLSF-estimated fluorescence lifetime map overlaid on the phantom. The regions of interest, left (L) and right (R), were selected using the intensity thresholding method. (b) Comparison of lifetime estimates for L and R using NLSF and single-snapshot RLD. (c) The leftmost and rightmost columns show continuous time-series single-snapshot acquisitions and the corresponding lifetime computations, respectively, while the phantom was in continuous motion (both positional and angular changes). Two random frames, frame 65 (green boundary) and frame 106 (red boundary), are enlarged in the second and third columns to display details of the intensity map and lifetime map, respectively.

Complete time-resolved fluorescence decay data were also captured for ground truth validation with traditional FLI using NLSF. The NLSF fluorescence lifetime map overlay (Figure 2(a)) illustrates the variation in lifetime values in two distinct areas of the embedding. The same phantom was subsequently imaged using the single-snapshot RLD-based method. Figure 2(c) leftmost and rightmost columns display the time-series single-snapshot data acquisition and its corresponding real-time fluorescence lifetime computation, respectively. During continuous single-snapshot data acquisition, the mouse-shaped phantom was rotated in various spatial positions and orientations. Figure 2(c) second and third columns highlight randomly selected frames from the time-series acquisition showing the positional change of the sample and overlaid intensity and lifetime computation. The time-series single-snapshot data acquisition and corresponding real-time FLI maps computation are shown in **Supplementary Video 2**.

Lifetime values estimated using NLSF and single-snapshot RLD-based methods are compared in Figure 2(b). The tube embedded on the left (L) contains AF700-PBS, while the tube on the right (R) contains AF700-DMSO. For the left embedding, both methods yielded similar average lifetimes, with the single-snapshot method exhibiting a larger standard deviation. In contrast, the right embedding showed both different average lifetimes and large standard deviations from the single-snapshot RLD method. The variation in lifetime estimation between the two methods is likely due to the phantom’s uneven surface profile. Differences in the topographical surface profile lead to pixel-wise variations in photon time-of-flight, causing offsets in the fluorescence decay profile. While the NLSF method accounts for these pixel-wise offsets when the IRF is used for re-convolution, the single-snapshot method estimates lifetimes without such offset corrections. In NLSF, lifetime is estimated using the excitation signal from the phantom (see *I*(*t*) in Equation 5).

Nevertheless, the single-snapshot RLD-based method provided sufficient contrast maps to distinguish the two distinct fluorescence lifetimes, effectively capturing their differing micro-environments in diffuse FLI. In contrast, the fluorescence intensity image (INT), while accurately locating the position of the embedded tubes, was not sufficient on its own to characterize differences in the fluorophores’ microenvironment.

To evaluate the applicability of wide-field FLI for surgical guidance, we performed a mock surgical procedure using a non-fluorescent mouse-shaped tissue-mimicking phantom with fluorescent embeddings (Figure 3) and a more complex fluorescent tissue-mimicking phantom with fluorescent embedding (Figure 4). These experiments aimed to demonstrate the potential of real-time FLI to assist in accurate localization and removal of distinct fluorophore embeddings based on their different lifetimes. [17]

**Fig. 3:**
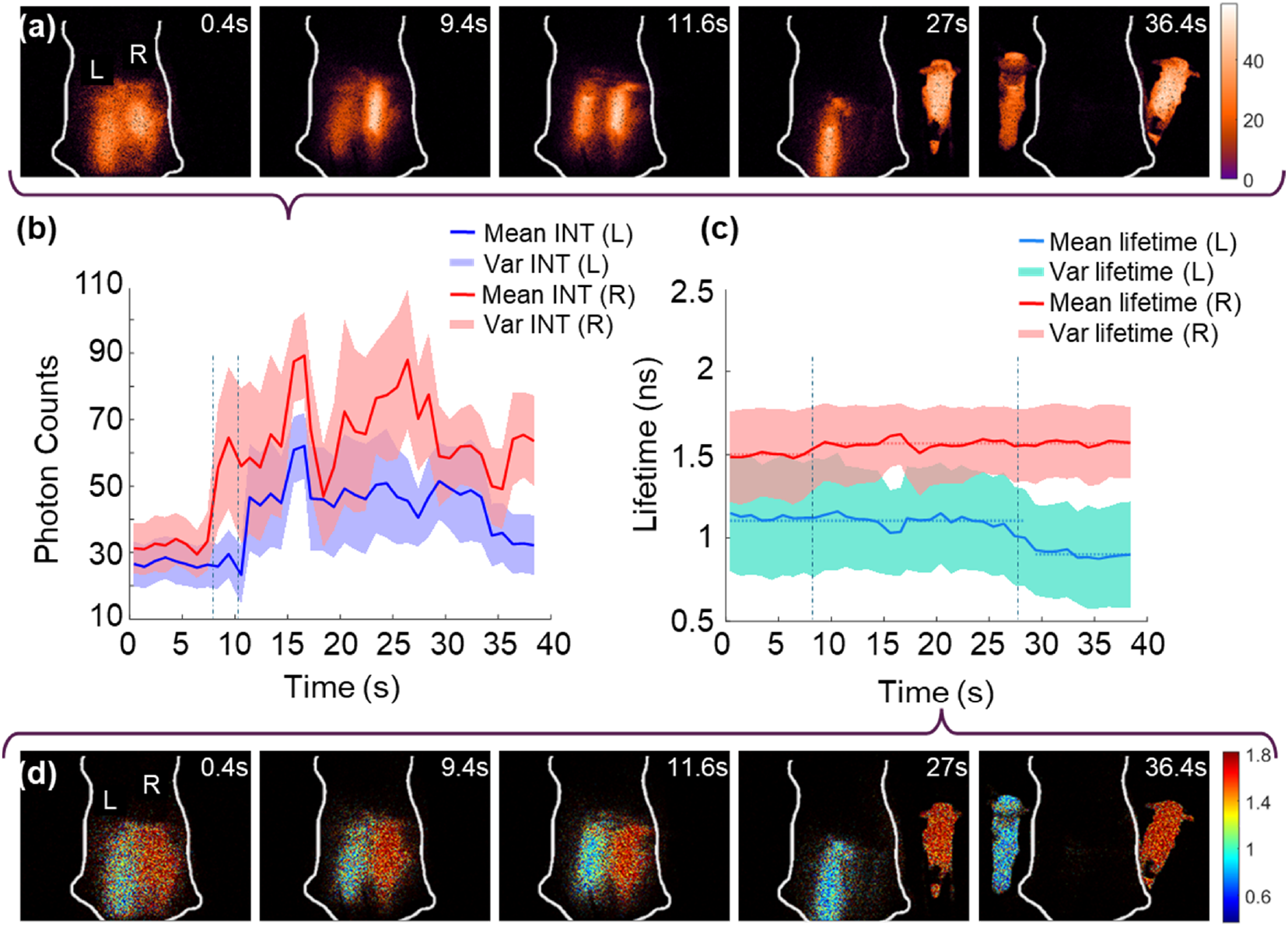
Real-time Macroscopic Fluorescence Lifetime-Guided Mock Surgical Procedure: (a) and (d) Time-series single-snapshot photon acquisition and real-time lifetime map computation, respectively. Both rows (left to right) show a mock surgical procedure performed on a tissue-mimicking, mouse-shaped phantom with fluorescence-embedded regions on the left (L) and right (R). The tubes were exposed by sequentially removing the upper layer (R first, followed by L) and were subsequently extracted from the phantom (R first, followed by L). (b) Photon counts of L and R embeddings with time. (c) Single-snapshot RLD-estimated fluorescence lifetime of the L and R embeddings over time. The white scale bar represents 10 mm.

**Fig. 4:**
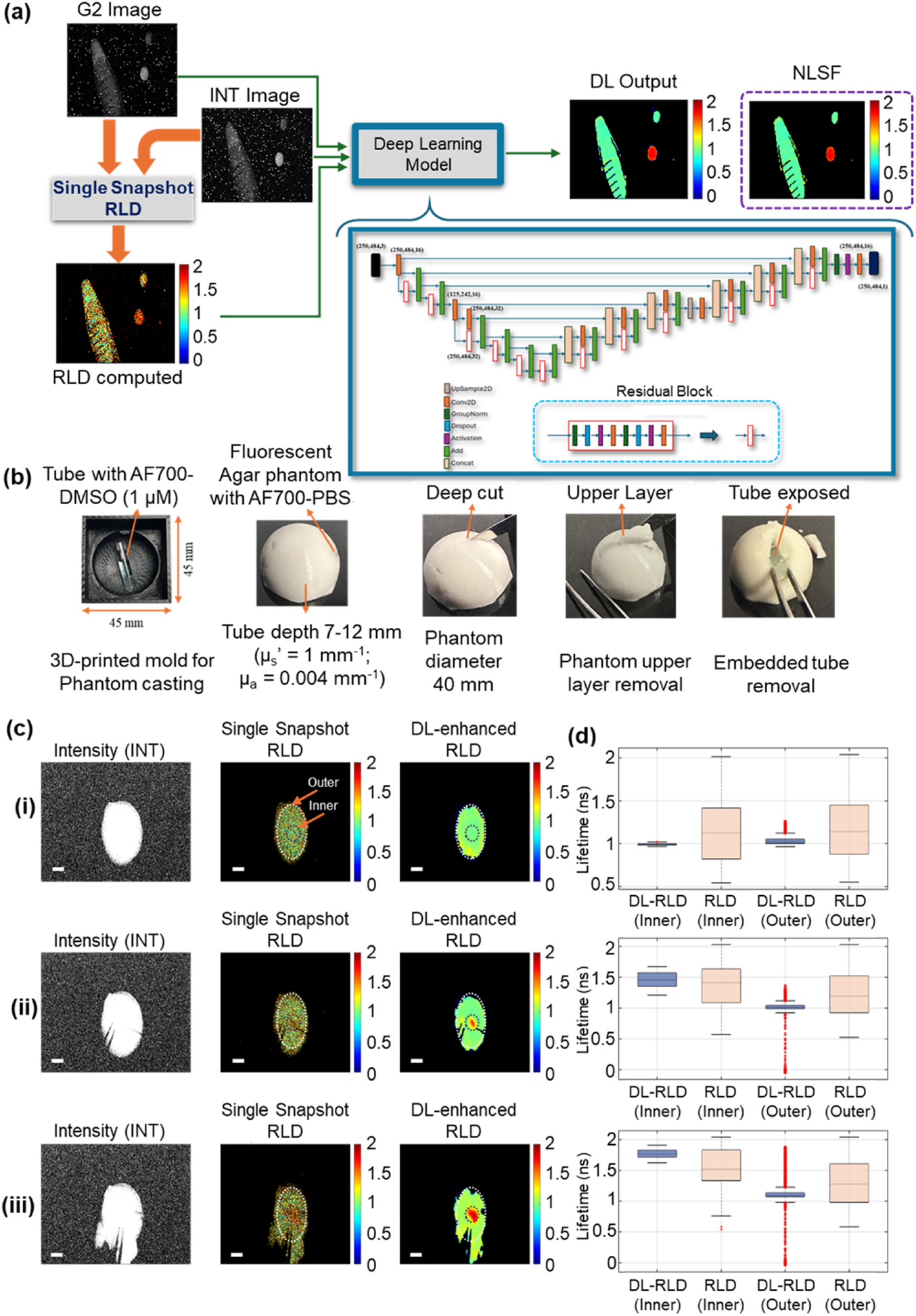
Deep Learning-Enhanced FLI for Guided Surgery in Fluorescent Tissue Phantoms: (a) Schematic representation of the DLenhanced single-snapshot RLD FLI workflow, illustrating the fluorescence lifetime estimation process using G2 and INT images processed with the RLD algorithm. The computed lifetime, G2 image, and INT image serve as inputs to the deep learning model (architecture shown in the inset), which generates DL-enhanced lifetime maps. The DL-enhanced result (output) compared with the NLSF estimated lifetime. (b) Preparation of a complex tissue-mimicking phantom, representing a human breast model, created using a 3D-printed mold. The phantom consists of a fluorescent base with fluorescent embeddings. The mock surgical procedure and embedding removal (left to right). (c) (i–iii) Three selected frames from the full time-series: (i) single-snapshot acquisition (INT), (ii) real-time fluorescence lifetime computation (singlesnapshot RLD), and (iii) corresponding DL-enhanced lifetime maps. The white scale bar represents 10 mm. (d) Single-snapshot RLD and DL-enhanced lifetime comparison for each frame (i–iii), using two regions of interest, inner (fluorescent phantom with embedding) and outer (only fluorescent phantom).

The videos of the real-time FLI-guided mock procedures and embedding removal are available in **Supplementary Videos 3 and 4**. Key time frames from **Supplementary Video 3** are shown in Figure 3(a) and (d), highlighting the single-snapshot photon count acquisition and real-time lifetime map computation, respectively. Each row displays intensity (top) and lifetime (bottom), while the sequence from left to right illustrates the systematic steps of the mock surgical procedure performed on the mouse-shaped tissue-mimicking phantom. The upper layers of the embedded regions on the right (R) and left (L) were removed at 9.4 s and 11.6 s, respectively, exposing the fluorescent embedding to the SS3 detector. Figure 3(b) shows the corresponding peaks in photon counts at these time points. Apart from these peaks, large variations in photon counts were observed throughout the surgical procedure; however, they consistently remained higher than the diffuse fluorescence photon counts.

In contrast to photon counts, the fluorescence lifetime values remained relatively stable throughout the procedure, as shown in Figure 3(c). The tube in region R was removed at 27 s, followed by the removal of the second tube at 36 s. A 50 ps lifetime difference was observed in tube R from the embedded to the exposed state. However, after this transition, the fluorescence lifetime of tube R remained nearly constant until the procedure ended with complete removal of the embeddings from the phantom. Tube L exhibited a consistently longer lifetime, by approximately 200 ps, while tube R remained inside the phantom. After the removal of tube R, tube L showed a consistent average lifetime of 0.9 ns, as highlighted at 27 s in Figure 3(c), which aligns well with the NLSF-estimated lifetime.

This experiment highlights the advantage of single-snapshot RLD in real-time FLI for the precise identification and removal of fluorescent embeddings, which is not feasible using fluorescence intensity data alone. Additionally, the experiment demonstrates real-time estimation of the impact of diffuse fluorescence signals from one fluorophore on another when they are in close proximity. The real-time FLI observed the sudden shift in fluorescence lifetime corresponding to an abrupt change in the surrounding environment.

### B. Single-Snapshot RLD Enhancement via Deep Learning

The signal-to-noise ratio (SNR) for a single pixel follows Poisson-limited temporal performance [18]. However, spatial uniformity, quantified as the SNR (average photon count divided by the standard deviation of photon counts across the array), falls below Poisson-limited SNR at low photon count levels due to non-uniform dark count rate (DCR) contributions, including hot pixels. As photon count levels increase, the relative impact of DCR non-uniformity diminishes, and the spatial SNR approaches the Poisson limit. At very high photon counts, sensor saturation reduces the standard deviation of counts below the Poisson statistics [18]. Dark count noise, pile-up effects, and temperature-dependent sensor characteristics also affect the raw single-snapshot data (G2 and INT channel) (see **Supplementary Section 1.1**). These effects can be experimentally characterized and numerically corrected [19] (see **Supplementary Section 1.2**). However, gate edge jitter, representing the stochastic temporal uncertainty in the gate edge position of individual pixels, and gate width variability remain the fundamental limitations that cannot be corrected and impact the SNR [18].

To address these challenges, we developed a deep learning (DL) model based on the widely used U-Net architecture [20]– [22], designed to increase the precision of single-snapshot RLD. The model architecture, illustrated in Figure 4(a), is fully described in **Supplementary Section 5**. The DL model can correct the artifacts introduced to raw data by the aforementioned multiple error sources. Moreover, it substantially minimized the pixel-wise variation in single-snapshot-based computed FLI maps. The DL-based enhancement of RLD showed higher accuracy benchmarked against NLSF estimated lifetimes obtained from full-decay acquisition, used as a reference due to the lack of a ground truth.

We evaluated the DL model’s performance using experimental data from a complex phantom that generated fluorescence signals throughout, including an embedded tube. This setup simulates a surgical scenario where target-bound probes have modified lifetimes, differentiating them from regions with passive probe accumulation and unchanged lifetimes [14]. In such cases, fluorescence intensity signals alone often fail to delineate embedding margins due to probe distribution throughout. In contrast, FLI can distinguish distinct tissue regions [23], based on environmental changes.

For this experiment, a tissue-mimicking phantom shaped to resemble human breast anatomy was prepared. The preparation steps for the phantom and bright-field images of the resection procedure are shown in Figure 4(b). The phantom was prepared by uniformly mixing AF700-PBS dye in the phantom matrix, ensuring that the entire phantom fluoresced under 700 nm pulsed illumination. To introduce a distinct fluorescence lifetime contrast, a cylindrical glass tube (inner radius 2 mm, outer radius 2.5 mm, and fill length ∼15 mm) containing AF700-DMSO was embedded at a depth of 7–12 mm beneath the curved surface of the phantom. Figure 4(c) depicts sequential steps of the embedding removal procedure. Figure 4(c) (i)-(iii) show, from left to right, the fluorescence intensity signal from the phantom, real-time single-snapshot RLD lifetime maps, and DL-enhanced fluorescence lifetime maps generated for the same frames. Figure 4(c) (i)-(iii) shows three randomly selected frames from the continuous acquisition during mock surgery of the complex phantom. The corresponding real-time experimental video is provided in **Supplementary Video 4**. The fluorescence lifetime map generated in real-time distinctly identified the location of the embedded tube, even before significant resection occurred.

As the procedure progressed, real-time FLI effectively guided the embedding region with each subsequent cut and guided the complete removal of the embedding, which exhibited a higher fluorescence lifetime than the background. Figure 4(d) (i)–(iii) demonstrate the DL-based enhancement in lifetime determination. For this analysis, the phantom was divided into inner and outer regions. The inner region was defined to encompass the entire embedding, while the remaining area was designated as the outer region. The DL-enhanced lifetime maps of the inner and outer regions showed significantly reduced pixel-wise variability compared to the single-snapshot RLD-based computation map. Both single-snapshot computed and DL-enhanced FLI methods provided comparable average lifetime estimates. However, the DL-enhanced FLI exhibited smaller pixel-wise variation and greater spatial uniformity.

### C. Real-Time Mesoscopic Volumetric FLI using Light-sheet Illumination

We applied the single-snapshot RLD method for fast 3D fluorescence lifetime reconstruction using a custom-built mesoscopic light-sheet fluorescence imaging setup (Figure 5(a)). For this experiment (see subsubsection IV-E.2), we imaged a breast cancer (AU565), HER2+ tumor spheroid treated with Trastuzumab (TZM) conjugated with NIR-I dye AF750 (TZM-AF750).

**Fig. 5:**
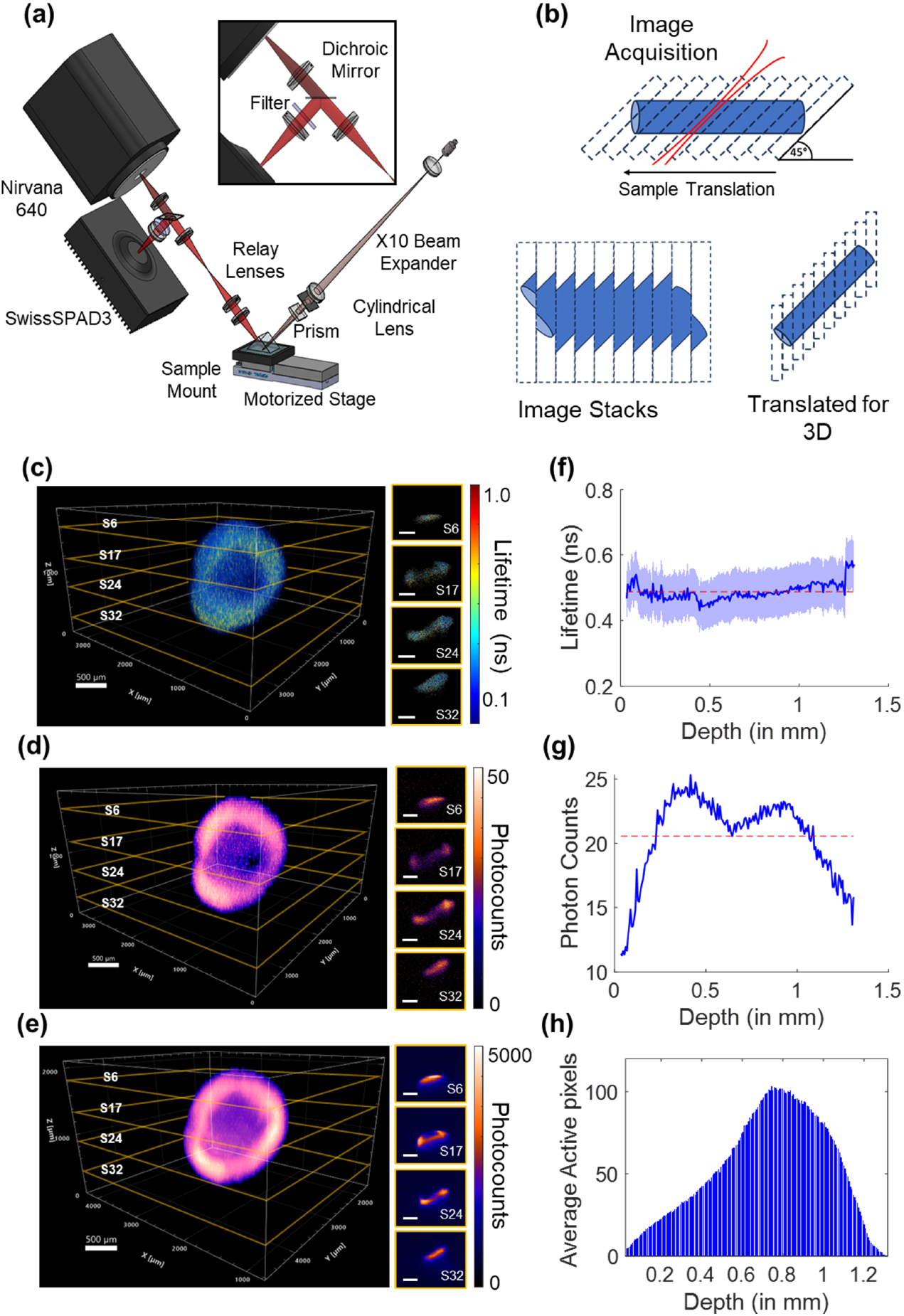
Mesoscopic Fast 3D FLI using Single-snapshot RLD: (a) Schematic of the mesoscopic fluorescence lifetime imaging (FLI) system employing light-sheet illumination and dual detection channels for near-infrared (NIR) and shortwave infrared (SWIR) fluorescence emission. (b) Schematic representation of angular image acquisition, stacking, and translational correction for 3D volume reconstruction. (c-e) 3D visualization of AU565 tumor spheroids treated with Trastuzumab-Alexa Fluor 750 (TZM-AF750); the white scale bars in 3D and 2D slices (s6, s17, s24 and s32) are 500 *µ*m. (c) Single-snapshot RLD 3D FLI reconstruction, (d) NIR-I 3D fluorescence intensity distribution derived from intensity (INT) images. (e) NIR-II fluorescence intensity maps acquired with an InGaAs detector, providing low-scattering 3D intensity reconstruction. (f) Single-snapshot RLD as a function of depth. (g) Average photon counts across tumor spheroid slices as a function of depth. (h) Average number of active pixels corresponding to tumor spheroid regions per slice as a function of depth.

This custom-built mesoscale light-sheet imaging set-up is designed to be equipped with two imaging modalities, SS3 for NIR-I FLI and an InGaAs detector for NIR-II fluorescence intensity imaging. Whilst the former is capable of single-snapshot real-time FLI, the latter enables deeper and higher resolution intensity-based imaging. Thanks to the high-quantum efficiency of InGaAs detectors, we can detect the long-tail emission of conventional NIR-I excited fluorophores, AF 750, in this experiment. Figure 5(b) shows a schematic of slice-wise imaging and its subsequent 3D reconstruction. The results of the mesoscopic light-sheet 3D imaging are summarized in Figure Figure 5(c)-(e). Single-snapshot RLD FLI slices for spheroid volumetric imaging are shown in Figure 5(c), the randomly selected slices S6, S17, S24 & S32 are shown to elaborate the illuminated part of a tumor spheroid in these slices and their corresponding real-time FLI computation. SS3’s corresponding intensity (INT) channel NIR-I images are shown in Figure 5(d). Figure 5(e) shows the NIR-II intensity images and volumetric reconstruction using the InGaAs detector. The 3D fluorescence intensity reconstruction in Figure 5(d) and (e) are from the INT channel of SS3 (NIR-I) and the InGaAs detector (NIR-II), respectively. Given the difference in pixel dimensions and sensor sizes between these two detectors, the images were cropped and resized for better tumor spheroid visualization. Figure 5(h) demonstrates the illuminated area of the tumor spheroid with depth, along with the average active pixels from which fluorescence signals were detected. While there was non-uniformity in the photon counts (Figure 5(g)) in the tumor spheroid imaging with depth, the single-snapshot RLD of the tumor spheroid remained mostly uniform, with slight variation in the middle (Figure 5(f)).

The FLI maps generated through single-snapshot RLD are shown in randomly selected slices in Figure 5(c), demonstrating fast volumetric mapping of fluorescence lifetime at the mesoscale. These results highlight the capabilities of the single-snapshot RLD method for rapid volumetric FLI at the mesoscopic scale. Light-sheet imaging techniques have previously facilitated fast, high-resolution fluorescence intensity volumetric imaging due to their sectioning ability. However, volumetric FLI has historically been constrained by lengthy time-resolved acquisition and processing requirements. By integrating the single-snapshot RLD method into a mesoscopic light-sheet imaging system, we successfully captured mesoscale volumes with thin sectioning in 2-3 minutes per imaging modality. This, therefore, enables fast mesoscopic mapping of probe biodistribution as well as fast retrieval of critical insights of the fluorophore microenvironment.

### D. Microscopic FLI: Calcium Imaging in Neuronal Cultures

To illustrate the capabilities of single-snapshot RLD FLI in the microscopic imaging regime, we used Cal-520^®^, AM, a dye emitting in the 520 nm range, to visualize calcium transients in murine cortical cultures over an 816 × *µ*m 816 × *µ*m field-of-view. The cells were submerged in Hank’s buffer with HEPES (HHBS) and maintained at 37°C with 5% CO_2_. Calcium activity was induced by application of 50 mM potassium chloride solution (KCl). Figure 6(a) shows the microscopy imaging setup, and Figure 6(b) depicts the capabilities of the full sensor (500 × 500 pixels) to capture high-resolution images across a large field-of-view. To fully demonstrate the dynamic single-shot RLD FLI ability of SS3, we focus on a single cell present in this sample by cropping down to a × 100 100-pixel region of the sensor (Figure 6(c)) and tracking the propagation of the intracellular calcium transient after chemical activation. By plotting the change in lifetime 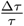 relative to an averaged baseline pre-activation as

**Fig. 6:**
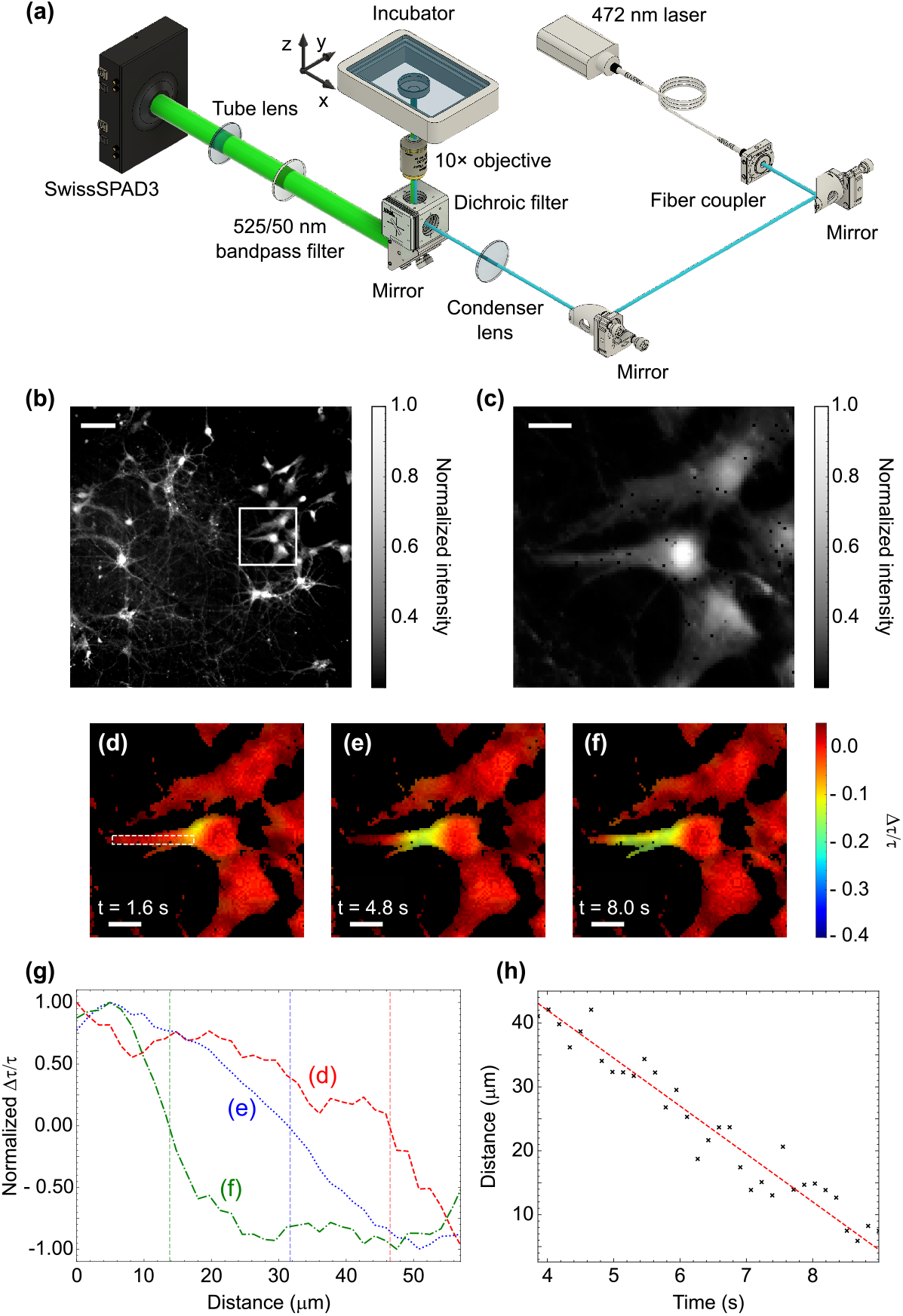
Microscopic Fluorescence Lifetime Imaging of Neuronal Calcium Transients: (a) Schematic of the microscopic fluorescence lifetime imaging setup. (b) 0.25-megapixel intensity image of neuronal culture. White scale bar, 100 *µ*m. (c) Cropped 100 *×* 100-pixel region enclosed in white border in (b). White scale bar, 25 *µ*m. (d) to (f) Frames from a 6.25-fps video acquisition showing the intensity-weighted lifetime change with the release of intracellular calcium within a glial cell following treatment with 50 mM potassium chloride solution (KCl). White scale bar, 25 *µ*m. (g) ∆*τ/τ* plotted along the cell body bounded by the white box (d) for frames (d) to (f). The halfway point in ∆*τ/τ* gives the wavefront position, with distance traveled indicated with vertical lines. Discretisation from the pixels was removed by interpolation. (h) Wavefront position versus time with associated linear fit giving a wavefront propagation speed of 7.47 ± 0.34 *µ*m/s.

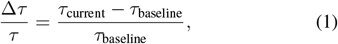

We can visualize the propagation of the calcium wavefront within the cell. The frames in Figure 6 (d) to (f) correspond to a ∼6.25 fps FLI acquisition and show a ∼15% change in lifetime where calcium ions are released. Real-time videos of 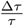 across the full field-of-view and of the cropped single cell are available in **Supplementary Video 5**.

We calculated the wavefront speed of the calcium transient by isolating a region-of-interest along the cell body aligned with the propagation direction (white box in Figure 6(d)). By plotting 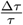 along the cell length, we can approximate the wavefront position for each frame in the video acquisition. Examples of this wavefront position for the frames given in Figure 6(d)-(f) are shown in Figure 6(g). From this analysis, and by linearly fitting through the resulting data (Figure 6(h)), we estimate the propagation velocity of the calcium wavefront to be 7.47 ± 0.34 *µ*m/s. This value agrees well with literature on fast intracellular calcium waves, [24], demonstrating this video-rate microscopic FLI technique to be a robust imaging modality for such dynamic interactions in active biological samples.

## III. Discussion

We have demonstrated the capability of single-snapshot-based real-time FLI using the SS3 system at ∼ 5 fps across diverse optical configurations (macro-, meso-, and micro-scale imaging) and in the visible to NIR spectrum of fluorescence signals. Such real-time FLI is particularly advantageous for time-evolving biomedical applications, where fast data acquisition and real-time analysis are critical.

Applying this method in wide-field diffuse fluorescence imaging, we closely mimicked *in vivo* animal imaging conditions, using tissue-mimicking non-fluorescent mouse-shaped and fluorescent human breast-shaped phantoms embedded with fluorescent samples. In the mouse-shaped phantom, the two embedded fluorescent samples contained the same fluorophore in different solvents, generating distinct fluorescence lifetime properties. The real-time FLI method successfully computed lifetime contrast maps, enabling clear visualization of distinct regions, a capability unachievable with conventional intensity-based imaging. The ability to differentiate spatially distinct regions in wide-field macroscale imaging using realtime FLI holds significant advantages for preclinical cancer imaging, drug kinetics studies (e.g., rapid drug internalization and clearance) [25], and real-time monitoring of dynamic processes.

Our method holds potential for clinical applications, such as fluorescence-guided surgery, by enhancing the identification of specific fluorescent markers based on their microenvironmental conditions, improving tumor margin detection, and facilitating their selective removal. We conducted a mock-surgery experiment using a complex tissue-mimicking breast-shaped phantom to simulate non-specific fluorescent probe accumulation, a common challenge in fluorescence-guided surgery. While traditional intensity-based imaging produced a uniform fluorescent signal across the tissue, real-time FLI effectively distinguished regions through fluorescence lifetime variations. Furthermore, DL-based enhancement of RLD, especially in challenging imaging conditions exhibited by complex fluorescent phantoms, further strengthens the clinical applicability of our method for real-time FLI-guided applications.

To extend real-time FLI beyond macroscale applications, we demonstrated its efficacy in mesoscale volumetric imaging of tumor spheroid models. Imaging at this scale provides an optimal balance between field-of-view and spatial resolution, making it well-suited for preclinical drug development research. By leveraging a mesoscopic light-sheet imaging system, realtime FLI enabled the rapid generation of volumetric fluorescence lifetime distributions. This capability is particularly critical for understanding tumor spheroid heterogeneity, which is invaluable in preclinical studies focused on drug response and resistance mechanisms. Moreover, fast volumetric FLI is essential for longitudinal studies where rapid imaging is crucial for capturing dynamic changes in tumor biology over time.

Finally, at the microscopic scale, our approach effectively captured rapid, dynamic cellular events. By resolving intracellular calcium transients in chemically stimulated neuronal cultures, we demonstrated the ability of single-snapshot FLI acquisition and the high spatial resolution of the SPAD camera to enable precise isolation and quantitative analysis of ion diffusion within individual cells across a large field-of-view. These findings highlight the potential of real-time FLI for high-resolution, dynamic cellular imaging. While our primary demonstration focused on calcium imaging, this approach can be readily extended to other fast biological processes, such as neuronal action potentials and cardiac conduction waves. With further enhancements in acquisition speed and field-of-view, this method could play a critical role in investigating a broad range of rapid cellular and tissue-level dynamics.

Beyond its technical capabilities, the real-time FLI method presented here offers substantial operational advantages. Even if the RLD methodology can, in principle, be achieved using a two-camera setup, such as an ICCD/ICMOS or a gated-SPAD camera paired with a standard co-registered CMOS/CCD camera (spatially and temporally synchronized) [26], [27], this dual detector approach would be bulky and cost-inefficient. But most importantly, such a 2-camera implementation would require absolute intensity calibration of both detectors across the entire data acquisition dynamic range for accurate RLD, which is extremely difficult to achieve and maintain due to its dependence on acquisition parameters that can vary (e.g., wavelength, high voltage, integration time, spatial nonlinearities, etc.). Furthermore, fast FLI data acquisition is often characterized by low photon counts, and different technologies exhibit varying noise characteristics, potentially biasing the results. As demonstrated in the macroscopic cases, the combination of SS3 and RLD provides robust results even when intensities fluctuate greatly due to motion/changes in locations. This system is inherently flexible, requiring minimal calibration and functioning effectively as a plug-and-play solution. Its ease of use, combined with fast data capture and real-time computation, makes it a powerful tool for both research and clinical applications.

Despite the advantages of the proposed approach, challenges persist due to time-of-flight (path length) variations, which result from changes in the height of the imaged object. In the case of the light-sheet mesoscopic system, time delays were accounted for based on the known geometry of the illumination light sheet (at a 45° angle). If these delays are not properly considered, they can cause a systematic shift in lifetime quantification with respect to depth (in the case of the mesoscopic system, ∼ 0.1 ns over 1.5 mm, see **Supplementary Figure 5**). However, in clinical applications, it may be difficult to account for these time shifts, as they could be linked to biological processes (breathing, motions, etc.) or external manipulation by an operator (e.g., a surgeon). Nevertheless, as shown in Figure 3(c), such biases are not significant when using lifetime-based contrast. Consequently, lifetime-based contrast is expected to remain useful and effective overall.

Another important aspect of the proposed methods is that their performance and imaging speed are intrinsically limited by the brightness of the sample. The speed of acquisition reported herein was normalized to 5 fps across all cases. This acquisition and processing speed were selected as it provided robust results for all experimental conditions. Though overall, it was constrained by photon-starved conditions in NIR applications. In these cases, higher frame rates are expected to be achieved based on readily available technological improvements. Especially, the SS3 units used in the mesoscopic and macroscopic regimes were not equipped with microlenses. Microlenses can enhance photon collection efficiency greatly by increasing the effective fill-factor to 84% [28], which can increase the NIR-I probe data acquisition speed sevenfold.

Likewise, higher speeds are anticipated in brighter samples as increased photon counts improve signal-to-noise ratios, in turn enabling faster data acquisition. Herein, the spectral range of the application plays a significant role in determining brightness and photon collection efficiency. In this study, we conducted microscopic FLI in the visible range, while mesoscopic and macroscopic FLI were performed in the near-infrared (NIR) spectral range. The latter is a far more challenging scenario due to three main factors. First, the quantum efficiency (QE) of SS3 is substantially lower in the NIR (10-20%; 700-800 nm) range compared to the visible range (35-50%; 500-650 nm range). Second, the brightness of visible fluorophores (quantum yield, QY ∼ 0.75-0.90%) is significantly higher than that of NIR fluorophores (QY ∼ 0.1-0.25%). Finally, the fluorescence lifetimes in the NIR (*<* ∼ 1 ns) are much shorter than those in the visible range (*>*∼ 2 ns), making variations in time delays more impactful and making late gates more susceptible to noise. As the primary limiting factor being the SNR, significantly improved frame rates should be achievable in microscopic settings due to the more favorable conditions (a couple of orders of magnitude).

Another key aspect of the work presented here is the real-time computational processes applied in the macroscopic application. For translational use, quantitative FLI readouts must be provided in real-time to the operator. In this work, on-the-fly data transfer and processing were executed using research-grade software environments such as LabVIEW and MATLAB. These processes accounted for ∼ 40% of the time required to generate a single frame. However, this time can be significantly reduced through more efficient coding practices and the integration of dedicated hardware acceleration techniques. In particular, leveraging AI and edge computing can address these bottlenecks, facilitating faster data processing and real-time analysis. As a first step, we utilized a U-Net model for inference, which is capable of delivering predictions within milliseconds. The model was specifically trained to handle noise and uncertainty, greatly enhancing the accuracy of FLI lifetime inference. However, it was deployed on a personal computer, necessitating data transfer and thus imposing some limitations on processing speed. Recently, we have reported the development of a neural network accelerator designed for on-board implementation (FPGA), specifically tailored to address hardware bottlenecks [9], [29]–[31]. The adaptation of these new edge computing models to RLD holds promise for significantly improving imaging speeds, enhancing user experience, and offering a more compact form factor. This progress makes bedside implementation not only more efficient but also more practical for clinical settings.

Lastly, the current implementation of RLD is limited to mono-exponential lifetime inference. In cases with more complex decay profiles, our approach can only infer the effective mean lifetime. Since bi-exponential decay models are common in many biomedical applications, modifications to the current method will be necessary. These might include interleaved gate acquisition [32], utilizing prior information about the single-exponential components of the decay, or further electronic advancements to enable simultaneous acquisition of at least one additional gate, while maintaining comparable data transfer rates. These technical challenges are expected to be addressed in the near future.

## IV.Methods

### A. SwissSPAD3

**Supplementary Section 1** provides detailed information on the SwissSPAD3 camera and its characterization, while the dual-gate signal acquisition architecture is illustrated in Figure 1(b). Following a frame reset signal, a 1-bit frame accumulation begins during which the SPAD is active (**Supplementary Fig. 1**). Within each frame, a user-defined number *N* of consecutive laser triggers are detected and used to generate a gate signal with a constant delay ∆*t* and duration *W*. A photon detected during *any of these gate periods* increments the gated signal 1-bit counter (G2) by one. A photon detected during *any of these laser periods* increments the intensity signal 1-bit counter (INT) by one. As a result, the output (INT, G2) of each frame can be either (0, 0), (1, 0), or (1, 1) (but not (0, 1)). A user-selected number of 1-bit frames *F* (F = 255 or 4 × 255) is then accumulated on the FPGA to obtain two images: the intensity image (INT) and the gated image (G2). In the case of a single-shot acquisition, this sequence is reproduced as long as needed. In the case of a fluorescence decay acquisition, the delay ∆*t* is incremented by a multiple of the gate step resolution (17.6 ps) after each set of (INT, G2) images. From the pair of images (INT, G2), the complementary gate *G*1 is trivially obtained by

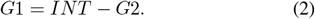

The gate signals have a rise and fall time of 200-300 ps. They exhibit a timing jitter of 109 ps and 153 ps FWHM for the rising and falling edges, respectively. The current implementation allows for frame rates (one INT and one G2 1-bit frame) of up to 49.8 kilo-fps. For more detailed characterization of the timing properties of the gates, see [12].

### B. Fluorescence Lifetime Decay Modeling

Fluorescence decay information can be captured using mainly two methods: frequency modulation and pulsed excitation. The pulsed excitation approach, also known as the time-domain approach, depends on the time-resolved recording of the fluorescence emission signal. In the simplest cases, the fluorescence decay (probability of detecting a photon at a time *t* after absorption of an excitation photon) follows a singleexponential law,

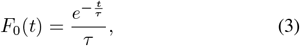

where *τ* is the fluorescence lifetime. The fluorescent sample is excited using a pulsed laser source with a single-pulse temporal profile *I*_0_(*t*), generating the incident fluorescence signal,

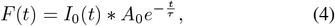

where *A*_0_ is the signal’s amplitude. This incident signal is detected by the SPAD with the single-photon electronic response function *E*_*SP AD*_(*t*) incorporating jitter, walk and other effects, adding uncertainty on the photon arrival time and detection efficiency *ϵ*, resulting in the detected signal *S*(*t*),

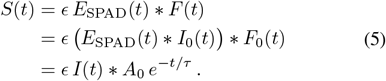

where *I*(*t*) = *E*_*SP AD*_(*t*) ∗ *I*_0_(*t*) is the global instrument response function (IRF). The T-periodic fluorescence decay *S*_*T*_ (*t*) captured by the SPAD is the infinite sum of single-pulse responses (Equation 5), offset by multiples of the laser period [33]. *S*_*T*_ (*t*) can also be written, up to a scaling factor, as the cyclic convolution of the T-periodic versions of both IRF and fluorescence decay (**Supplementary Section 2.1** and ref. [33]).

Acquisition of the temporal decay of a sample using time-gated detectors is time-consuming and depends on multiple factors such as gate image exposure time, delay between consecutive gates, and the number of gates needed to capture a decay curve. Long acquisition times prohibit FLI application in dynamic imaging scenarios, such as studying rapidly evolving biological processes, monitoring living organisms, and clinical applications like fluorescence lifetime-based tumor resection and guided surgery. Additionally, very long acquisition increases the risk of photobleaching, further constraining its utility.

### C. Rapid Lifetime Determination

The RLD approach is a computationally efficient method for the very fast estimation of fluorescence lifetimes [34]–[38]. It constitutes a class of algorithms applicable to both single- and multi-exponential decays, with or without a baseline offset. Unlike traditional methods that rely on full decay curve fitting, RLD utilizes time-integrated intensity signals over specific intervals of the decay curve to algebraically calculate the lifetime − significantly reducing computational complexity. Time-gated cameras (SPADs & ICCDs) serve as ideal platforms due to their inherent ability to integrate photon counts over predefined time intervals. A brief discussion of the underlying principles and implementation strategies is provided in **Supplementary Section 2**. Conventional RLD approaches typically require multiple contiguous or overlapping acquisitions along the decay curve to compute lifetimes. However, with the novel dual-channel capability of the SS3 detector, precise lifetime estimation is possible using only a single acquisition, referred to as single-snapshot acquisition. This approach eliminates the need for iterative or sequential measurements, thereby paving the way for real-time lifetime determination with improved accuracy and efficiency.

#### 1) Fast lifetime computation with SwissSPAD3

The temporal profile of fluorescence emission can often be modeled as a mono-exponential decay, as described in Equation 3. Traditionally, fluorescence decays recorded by time-resolved detectors are subsequently analyzed using methods such as NLSF, maximum likelihood estimation (MLE), or Bayesian approaches to determine the fluorescence lifetime parameters. These approaches are computationally intensive and time-consuming. Alternatively, methods based on projections of the decay on an orthogonal basis of functions (sine and cosine in phasor analysis [39], [40], Leguerre polynomials [41], etc.) have been proposed, which reduce the computational burden but, in general, require the recording of complete decays to be used.

In contrast, RLD algorithms utilize contiguous or overlapping gated acquisitions, and (using a few assumptions about the decay) algebraically compute the lifetime. The RLD method can be naturally adapted to leverage the dual-gate acquisition mechanism of the SS3 system, allowing fluorescence lifetime estimation from a single-snapshot acquisition. The following details the relevant equations needed for this analysis, using notations from ref. [33].

##### a) Time-gated periodic decays: Excitation pulse, pure decay, and emitted signal

Consider a *T* -period pulsed laser, as depicted in the schematic Figure 1(b). At steady state, the *T* - periodic excitation of the system results in a *T* -periodic emitted signal, with temporal profile equal to the sum of the signals excited by each individual laser pulse. Assuming the sample is excited by a Dirac delta pulse *δ*(*t*), the resulting emission is denoted as *F*_0_(*t*), which represents the pure decay response of the fluorescence sample as described in Equation 3. In practical systems, the pulsed excitation is not instantaneous and has a finite width temporal profile, *x*_0_(*t*), which results in an emission signal, *ε*_0_(*t*) expressed as the convolution product,

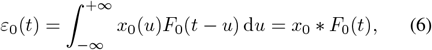

where *F*_0_(*t*) is equal to zero for *t <* 0 and decays from a maximum value reached at *t*_max_ ≥0 to 0 as *t*→ ∞.

##### b) T-periodic summation and periodic signal

The steadystate *T* -periodic emission signals *ε*_0,*T*_ (*t*) obtained by the summation of the responses to infinitely many excitation pulses separated by a period *T* are given by the *T* -periodic summation

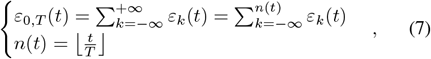

where ⌊*x*⌋ is the largest signed integer *n*≤ *x* (the sum truncation in Eq. 7 simply reflects the fact that excitations taking place after time *t* cannot contribute to the signal at *t*) and the infinite series of excitation pulses *x*_*k*_(*t*) and single-pulse emission decays *ϵ*_*k*_(*t*), indexed by the signed integer *k*, are defined by

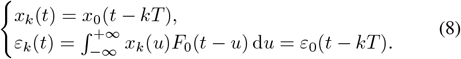

The index *T* in *ε*_0,*T*_ in Equation 7 indicates that it is a *T* periodic function, as will be the convention in the remainder of this article.

##### c) Instrument response function

The emitted signals are recorded using multiple instruments (detectors, electronics, etc.) which have a characteristic response *E*(*t*) to an instantaneous signal *δ*(*t*) (e.g., a single photon). The recorded signals from the *T* -periodic emitted signal can be written as a convolution of periodic *ϵ*_0,*T*_ and non-periodic *E*(*t*).

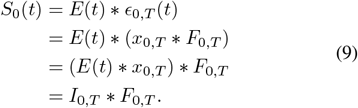

Equation 9 introduces the *T* -periodic instrument response function *I*_0,*T*_. Hence, the recorded signal will be the cyclic convolution of the IRF and periodic sample decay and can be written as

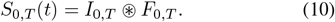

##### d) SwissSPAD3 dual gating

If *s* represents the gate offset relative to a reference trigger, typically offset with respect to the excitation pulse, and *W* denotes the gate width, the gating function of SS3 can be approximated as an ideal square (boxcar) function. For simplicity, the non-instantaneous response of the detector to the applied voltage is neglected. This response introduces minor deviations from the ideal square shape due to voltage transients; however, their impact on the overall analysis is negligible and can be reasonably ignored.

The gates are synchronized with the excitation pulse, and data are captured at each period *T*. Consequently, the periodic version of the gating function can be expressed as

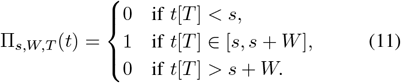

For single-snapshot acquisition, the values of *s* and *W* are kept constant for each trigger period. Assuming that *s* + *W* ∈ [*W, T*], the functions for the G2 and INT signals can be defined as follows:

- **G2 Gate** (II(G2)): for constant s and W.

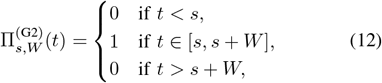

- **INT Gate** (II^(INT)^): where s = 0 and W = T.

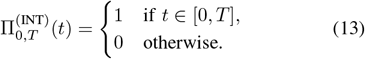

The signal accumulated during a square gate starting at time s, *S*_*T,W*_ (*s*), is an integral of the product of the square gate with the signal *S*_*T*_ (*t*),

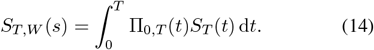

The accumulated G2 and INT signals can be represented as 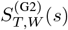 and 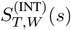

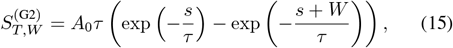

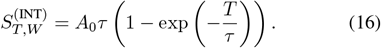

The accumulated signal ratio *R*_*T,W*_ (*s*) is the ratio of the accumulated G2 signal to the accumulated INT signal,

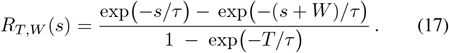

and in case of *s* = 0, the Equation 17 can be reduced to

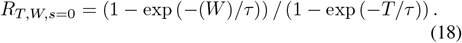

Because this equation is not algebraically invertible to obtain *τ* from *R*_*T,W,s*=0_, a precomputed lookup table relating *τ* to the observed *R*_*T,W,s*=0_ can be used to solve for Equation 18. The details of the pixel-wise IRF offset correction and look-up table for *R*_*T,W,s*=0_ are explained in Supplementary Section 2. A series of experiments were conducted in the micro-, meso, and macro-scale FLI setups to compare RLD-based lifetime estimation with the conventional lifetime estimation methods. It has been shown that this RLD methods with the SS3 camera enable the reconstruction of lifetime images with ∼ 1/100th less acquisition time compared to full temporal data capture in the same SPAD camera.

### D. Sample Preparation

#### 1) Primary Neuronal Culture Preparation

Cortices were isolated from the brains of E18 embryos of C57BL/6 mice and collected in ice-cold HBSS buffer (Hanks’ Balanced Salt Solution). The cortical tissue was chopped into fine pieces, washed in HBBS, and incubated with TrypLE Select 10X for 10 minutes at 37^*°*^C. The TrypLE 10X was then inactivated by the addition of 7 mL Neurobasal Plus complete media (supplemented with B-27, glutaMAX, and penicillin/streptomycin) and centrifuged at 1200 rpm for 5 minutes at room temperature. The pellet was resuspended and subjected to mechanical agitation to break down any remaining clumps of cells. Cells were seeded at 1 × 10^6^ cells/mL onto a 6 well-plate with coverslips, precoated with poly-D-lysine and laminin (4 *µ*g/mL). The cultures were maintained in Neurobasal Plus complete medium at 37^*°*^C with 5% CO_2_. The animal work and care were carried out under a UK Home Office project license under the Animals (Scientific Procedures) Act (1986).

To perform calcium imaging, the 14 DIV cortical cultures were loaded with 5 *µ*M Cal-520^®^, AM for 50 minutes at 37^*°*^C and washed for another 20 minutes with Hanks’ and HEPES Buffer (HHBS, 20 mM HEPES) at 37^*°*^C. During the experiment, the coverslips were kept submerged in HHBS. Baseline measurements were recorded without any external stimulus. Stimulus was delivered in the form of extracellular application with 50 mM KCl. Calcium transients induced by the application of KCl were observed.

#### 2) Tissue Mimicking Phantom preparation

To prepare the phantom, we combine distilled water, 1% India Ink (Speedball Art Products, NC), 20% intralipid (Sigma–Aldrich, MO) of volumes of 157.05 mL, 1.05 mL and 11.90 mL, respectively with 1.7 g of agar to form a homogeneous phantom that has roughly the same background optical properties as the in silico phantoms used in training. Two appendroff’s size cavities are made in the phantom at a depth of 2-3 mm, with the centers ∼10 mm apart, in which Alexa Fluor 700 dissolved in PBS and DMSO, respectively, was placed carefully. The rest of the phantom was poured completely to encapsulate the entire shape.

#### 3) Liquid Overlay Tumor Spheroid Preparation

Human epidermal growth factor receptor 2 (HER2+) AU565 cells (ATCC CRL-2351; breast cancer) were cultured in complete RPMI 1640 cell culturing medium containing HEPES supplemented with fetal bovine serum and penicillin/streptomycin. Cells were washed once with warm PBS, incubated with TrypLE Express for 5 minutes at 37^*°*^C to detach them and resuspended in fresh complete RPMI medium at a concentration of 500,000 cells/mL. Matrigel was thawed over ice for an hour, then a 10% Matrigel media was made using complete RPMI. 50 *µ*L of 10% Matrigel media was pipetted into the wells of a low-cell adhesion 96-well plate. The AU565 cell suspension was flicked to resuspend settled cells, and then 50 *µ*L of cell suspension was pipetted into each well containing 10% Matrigel media. A counterbalance was made by pipetting 100 *µ*L of PBS into the respective wells of another low-adhesion plate. The plates were then centrifuged at 1000 RPM for 10 minutes. Spheroids were left to grow in a 5% CO_2_ incubator at 37^*°*^C for 4 days prior to treatment.

To label cells, a 60 *µ*g/mL solution of Trastuzumab (MedChemExpress HY-P9907) conjugated to AF750 NHS ester (ThermoFisher A20011) was prepared in complete RPMI media. Media was removed from the wells of the low adhesion plate and replaced with the Trastuzumab-AF750 containing media. No washes were performed. Spheroids were incubated for 22 hours prior to media removal. Spheroids were then rinsed with PBS and transferred to a dish containing phenolred free DMEM with HEPES for imaging.

### E. Multiscale Fluorescence Lifetime Imaging Set-ups

#### 1) Microscopic FLI set-up

The microscopic imaging system shown in Figure 6(a) was based on an inverted fluorescence microscope design. Briefly, excitation was provided by a HORIBA DeltaDiode operated at 80 MHz (DeltaDiode-L, 470 nm, HORIBA Scientific, Glasgow, Scotland), with a peak wavelength at 472 nm and a typical pulse width of 65 ps. This was launched through a 150 *µ*m^2^ diameter square-core multimode fiber (M101L02, Thorlabs, NJ, USA) to mitigate uneven illumination arising from a Gaussian-shaped intensity profile. We also removed any speckle patterning from the illumination by vibrating the fiber throughout experiments [42]. The square beam was focused onto the back aperture of a 10× objective (Nikon 10× /0.50 NA) with a 100 mm achromat (AC254-100-A-ML, Thorlabs, NJ, USA) to provide collimated excitation to the sample. The objective was mounted on a piezo-controlled mount (PFM450E, Thorlabs, NJ, USA), to improve control over image focus. Emission signal was collected and separated from excitation using a FITC emission/excitation filter and dichroic mirror (MD499, Thorlabs, NJ, USA) and filtered further with a 525/50 nm bandpass filter (ET525/50 M, Chroma Technology Corp., VT, USA). The resulting signal was imaged onto the SS3 sensor [12] with a 200 mm tube lens.

To ensure that the samples were maintained at physiological conditions, a heated incubator was mounted on the system above the objective. CO_2_ and air conditions were controlled with a two-gas mixer (2GF-MIXER, Okolabs, PA, USA) and delivered to the incubator containing the sample (H301-K-FRAME, Okolabs, PA, USA). We monitored the state of this incubator and controlled the temperature via the control panel connected (OKO-TOUCH, Okolabs, PA, USA). This incubator was mounted on a precision XY scanning stage (MLS203-1 controlled by a BBD202, Thorlabs, NJ, USA) to allow complete control over sample position.

To demonstrate this system’s RLD capabilities at the microscopic scale, we performed calcium imaging on a sample of 14 DIV cortical cultures, focusing specifically on the intracellular calcium transients in a glial cell. Sample preparation is outlined in section subsubsection IV-D.1. SS3 was set to capture 12-bit images consisting of 4096 consecutive 1-bit images captured with an exposure time of 30.72 *µ*s. The gate was triggered at 40 MHz by setting a 1/2 clock divider on the laser output trigger. We chose a 2 ns gate duration to match our predicted lifetime.

#### 2) Mesoscopic FLI set-up

We developed a light-sheet-based illumination setup for meso-scale FLI. Briefly, emission from the same tunable Ti: Sapphire laser as the Macroscopic FLI system was coupled into a 50 *µ*m core diameter multimode optical fiber (M14L10, Thorlabs, NJ, USA), where the output was connected to a collimator (F220SMA-780, Thorlabs, NJ, USA). To generate a light sheet, the output beam was conditioned, first through a beam expander made from two achromatic lenses (LA1951-B and LA1461-B, Thorlabs, NJ, USA), followed by truncation through adjustable mechanical slits (VA100CP, Thorlabs, NJ, USA), and finally focusing through a cylindrical lens (LJ1703RM-B, Thorlabs, NJ, USA), achieving a light sheet thickness of ∼ 50 *µ*m. The resulting light sheet was focused onto the sample at a 45-degree angle. We compensated for refractive index mismatch by both illuminating the sample and imaging through a right-angle prism (PS611, Thorlabs, NJ, USA), which was in contact with the sample [43]. Emission signal was collected and relayed through a series of achromatic lenses (MAP105050-B and AC254-040-B-ML, Thorlabs, NJ, USA) into a dichroic mirror (DMLP950R, Thorlabs, NJ, USA), which split the fluorescent signal into two channels at 950 nm. Signals at wavelengths below 950 nm were filtered using a 832 ± 37 nm bandpass filter (FF01-832/37, Semrock Inc, USA) and focused into a SwissSPAD3 detector through an achromatic lens (AC254-050-B-ML, Thorlabs, NJ, USA). Similarly, signals at wavelengths above 950 nm were filtered using a 1000 nm long pass filter (FELH1000, Thorlabs, NJ, USA) and focused into a liquid-cooled InGaAs camera (NIRVANA 640, Teledyne Princeton Instruments, NJ, USA) through an achromatic lens (AC254-050-C-ML, Thorlabs, NJ, USA). To achieve volumetric imaging, samples were translated through the excitation light sheet using a motorized translation stage (MTS50-Z8, Thorlabs, NJ, USA). Given the light sheet excitation at a 45-degree angle with respect to the direction of sample translation, we used a step size of 70.71 *µ*m to match the thickness of the light sheet.

To demonstrate this system’s RLD capabilities at the mesoscopic scale, we performed volumetric imaging of a tumor spheroid after treatment with Trastuzumab conjugated with NIR-I dye AF750 (ThermoFisher, USA). Sample preparation is outlined in subsubsection IV-D.3. The sample was mounted onto the imaging system through a custom 3D printed mount filled with cell media, and imaging was performed by translating the sample through the light sheet illumination. SWIR intensity images were acquired with an exposure time of 1 second per acquisition, where the detector was cooled to -80^*°*^C to minimize dark current noise. Similarly, NIR-I RLD images were captured with an exposure time of 20 ms per acquisition. Image processing, including shift correction resulting from the angled imaging for volume visualization, was performed using custom MATLAB scripts.

#### 3) Macroscopic FLI set-up

An illustration of the SwissSPAD3 (SS3) macroscopic FLI configuration is shown in Figure 1(a). The technical details about the SS3 camera can be found in an earlier publication [44] and are also explained briefly in subsection IV-A and **Supplementary Section 1**. A tunable Ti: Sapphire laser (Mai Tai HP, Spectra-Physics, CA, USA) with a laser repetition rate ∼ 80 MHz (*f*_laser_) was used as an excitation source. The laser excitation was directed to the sample plane using a multimode optical fiber (QP200-2-VIS-NIR, Ocean Optics, FL, USA). The wide-field illumination was projected onto the sample plane using a Digital Micromirror Device (D4110, Digital Light Innovations, TX). The emitted fluorescence signals were collected through an application-specific bandpass emission filter by a macroscopic photographic lens (AF Nikkor 50 mm f/1.8D, Nikon, Tokyo, Japan). SS3 was set to acquire 10-bit images consisting of 1020 accumulated 1-bit gate images, with each 1-bit image resulting from exposure of each SPAD pixel to the incoming photon flux for a user-specified duration [45]. The optical imaging is performed in reflective geometry with a field-of-view of ∼ 40 × 40 mm^2^.

## V.Acknowledgments

The authors would like to acknowledge the generous funding received from the National Institutes of Health under grants R01-CA271371, R01-CA237267, and R01-CA250636, and from the UK Engineering and Physical Sciences Research Council (grant numbers EP/T00097X/1, EP/T021020/1). DF acknowledges the Royal Academy of Engineering’s support through the Chairs in Emerging Technology Program.

## Notes

### Competing Interest Statement

The authors have declared no competing interest.

