## Supplementary material for "Real-Time Wide-Field Fluorescence Lifetime Imaging via Single-Snapshot Acquisition for Biomedical Applications": arxiv_singleshot_supplementary.pdf

Presented here are the supplementary details of the manuscript titled "Real-Time Wide-Field Fluorescence Lifetime Imaging via Single-Snapshot Acquisition for Biomedical Applications."

### I. SwissSPAD3 (SS3)

SwissSPAD3 (SS3) [1] is a 500×500-pixel single-photon avalanche diode (SPAD) camera, fabricated in 0.18 $\mu$m CMOS technology. Each pixel is composed of a frontside illuminated PIN photodiode with an active-area diameter of 6 $\mu$m Figure 1(a), tiled over the 500×500 array and a pixel pitch of 16.38 $\mu$m Figure 1(b). When compared with its predecessor, SwissSPAD2 [2], SS3 offers a novel dual-gate architecture, improved gate timing properties, and a smaller minimum gate window duration (1 ns), whilst maintaining excellent SPAD detection characteristics and a similar form factor Figure 1(d). The system is housed on a motherboard PCB and comprises the sensor, two Opal Kelly XEM7360 FPGA boards, and a microcontroller to configure various supply voltages.Figure 1(c) shows details on the sensor. The readout architecture of SS2 is based on a global shutter, while that of SS3 is based on a rolling shutter. This means that SS3's rows are exposed in synch with the light source but slightly delayed, 20 ns, from each other.

SS3 achieves a peak photon detection probability (PDP) of over 50% at 520 nm, as shown in Figure 1(e), and maintains a relatively high PDP over much of the visible spectrum. The fill factor of the sensor is 10.5%, but the addition of imprinted microlenses can increase this value to ≈ 50%, in a large visible spectrum, provided relatively high collimation of the main optical system [3]. The SPAD design affords low dark count rate (DCR) operation, as shown in Figure 1(f). At typical operating conditions with a $V_{excess}$ of 6 V, a median DCR of less than 10 cps is achieved.

An example of the raw data (INT and G2 channels) acquired in macroscopic FLI set-up is discussed in subsection I-A. Various factors, such as dark count noise, pile-up effects, and temperature-dependent sensor characteristics, introduce artifacts into the raw data (see subsection I-B). For our RLD algorithm implementation, these effects were experimentally characterized and numerically corrected.

#### A. SS3 Raw Data (without corrections)

As explained in the main paper **Section 4.1**, the INT and G2 images were captured continuously within a user-defined gate. An example of raw data from the diffuse signal from a white paper at 700 nm illumination and a fluorescence image using a tissue-mimicking phantom in the shape of a mouse is shown in Figure 2(a) and Figure 4(a), respectively. Figure 2(b) and (c) show time-resolved acquisition of the INT and G2 channels for four randomly selected pixels, highlighting the INT channel integrates the entire fluorescence signal, whereas the G2 channel focuses on a chosen temporal segment of the decay. Figure 2(d) shows the spatial photon count variations at randomly selected gate number 61.
Figure 2 (e-g) shows photon variation in time-series (multiple frames) data acquisition. Herein a time-resolved acquisition of a single pixel at (350,164) is compared across 4 consecutive time-series frames (n to n+4) where n is a randomly selected frame in multiple time-resolved acquisitions made in the same field-of-view. Figure 2(g) shows the temporal photon count variations at randomly selected gate number 61.
A similar single-snapshot raw data set is presented in Figure 4, where the same four-pixel comparison Figure 4(b–d) and temporal photon counts variations of single-pixel Figure 4(d–f) are shown in a tissue-mimicking phantom with AF700 in DMSO and AF700 in PBS embeddings.

#### B. SS3 Raw Data, Artifacts & Corrections

**Dark Counts** are thermally generated detections that occur in the absence of incident photons. In SS3, dark counts are

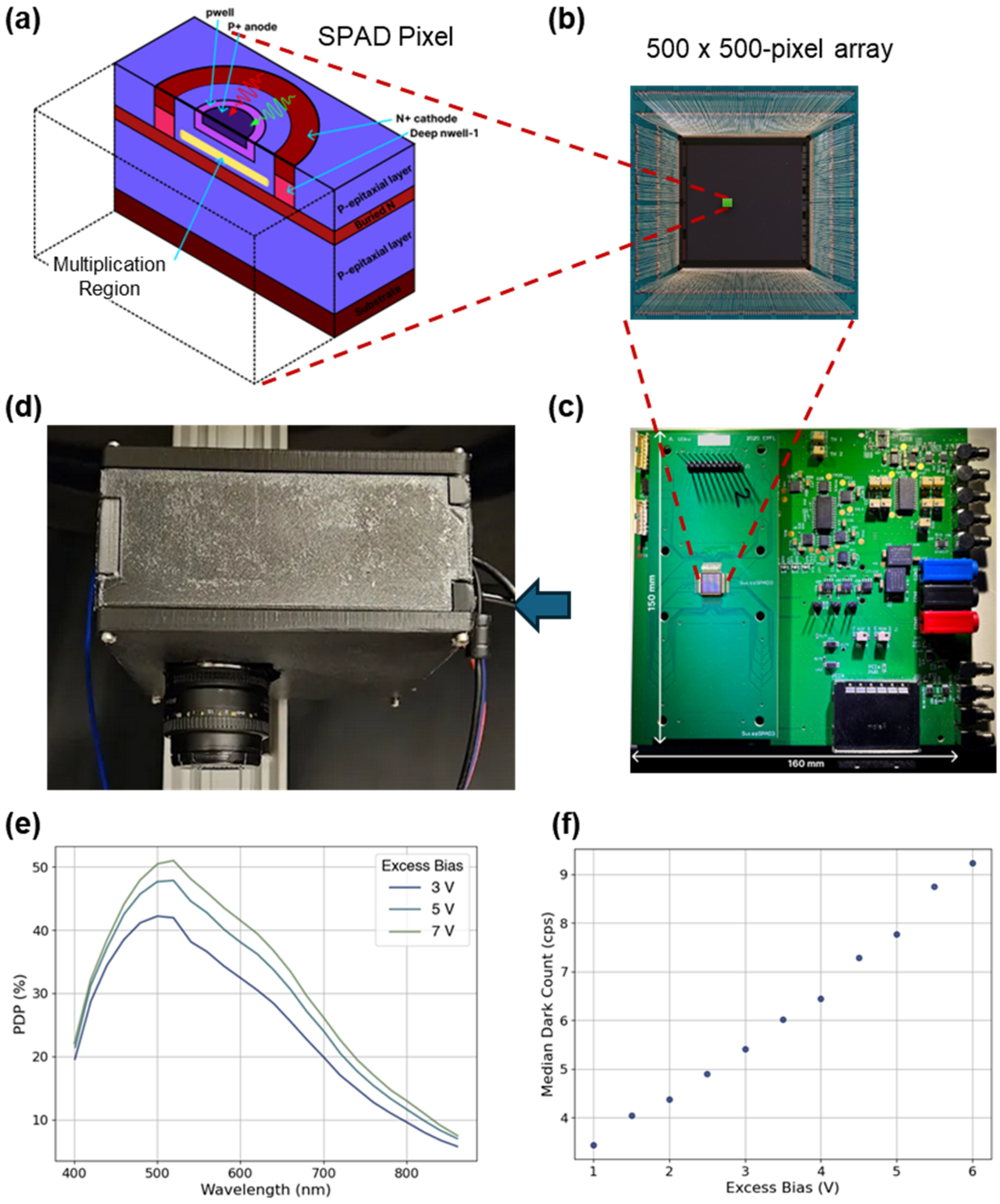


Fig. 1**: Time-gated SPAD detector SwissSPAD3 (SS3).** (a) Schematic of the SPAD pixel cross-section for p-i-n SPAD arrangement. (b) 500 $\times$ 500-pixel array for wide-field imaging. (c) SPAD sensor arrangement on the electronics board, including the FPGA and PCIe slots. (d) Assembled small footprint detector. (e) Photon detection probability (PDP) of SPAD array as a function of wavelength. (f) Median dark count rate as a function of excess bias voltage.

one of the primary sources of noise. In the SS3 detector, the dark count rate (DCR) is defined as the average number of dark counts per unit time. Dark counts follow a Poisson distribution, indistinguishable from counts caused by photons. The time interval between two adjacent dark counts on the same SPAD follows an exponential distribution, when far away

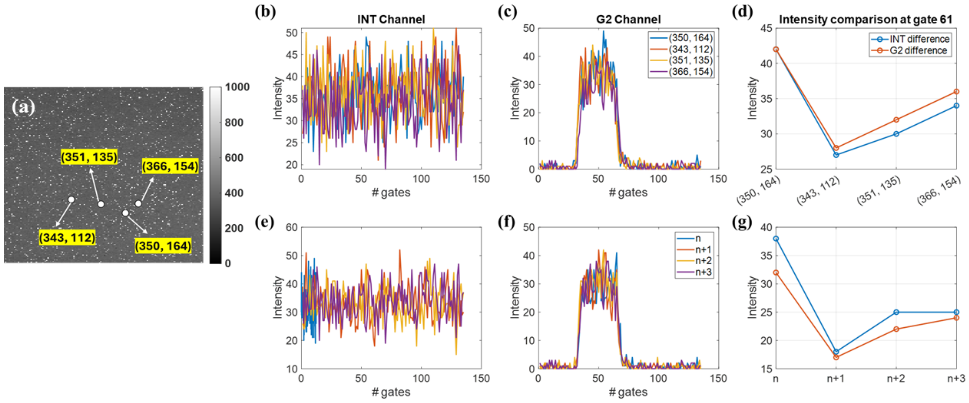


Fig. 2**: IRF intensity information** from (a) 4 random selected pixels of (b) INT channel, (c) G2 channel, and (d) corresponding intensity differences at gate 61, and intensity information from pixel (350, 164) at different imaging gates, n, for (e) INT channel, (f) G2 channel, and (g) corresponding intensity differences at gate 61

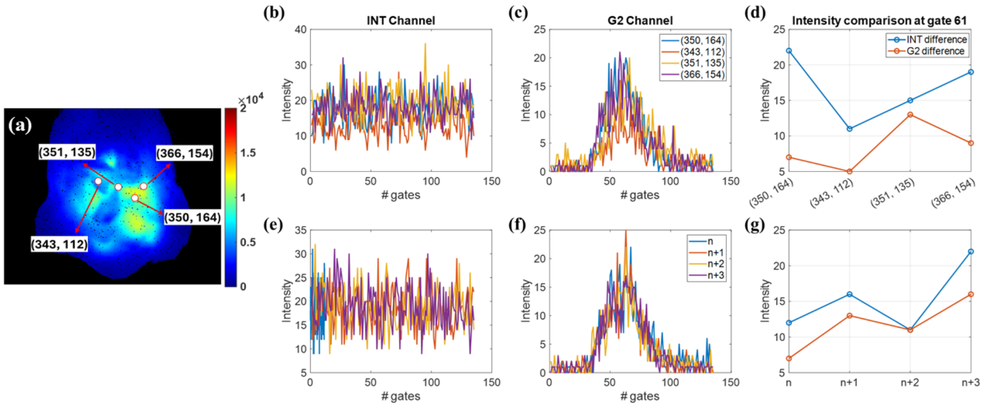


Fig. 3**: In Silico mouse fluorescence intensity information** from (a) 4 random selected pixel of (b) INT channel, (c) G2 channel, and (d) corresponding intensity differences at gate 61, and intensity information from pixel (350, 164) at different imaging gates, n, for (e) INT channel, (f) G2 channel, and (g) corresponding intensity differences at gate 61

from each other, otherwise afterpulsing may arise, since the quenching of the SPAD occurs upon an event and recharge when the next time gate starts. Over an array, the DCR distribution among SPADs can often be approximated by a normal distribution. The uniformity of DCR between pixels affects the spatial resolution of the image. DCR is typically quantified using a combination of the average DCR and the percentage of hot pixels in the array. Dark count correction can be implemented through multiple experimental observations, while the proportion of dark counts due to afterpulsing can be estimated but not completely suppressed. It is important to note that only one photon can be detected during the readout cycle. As a result, any count loss or **pile-up** must be corrected during post-processing. A correction formula from [4] is used for this purpose.

**Pixel Crosstalk** is another source of error that must be corrected in raw data. Pixel crosstalk occurs when detection events in one pixel are triggered by an event in an adjacent pixel, introducing unwanted correlated noise. Pixel crosstalk can electrical and optical. In SS3, bothe forms of crosstalk were minimized by use of deep trench isolation (DTI) and by careful design of the pixels. Using pixel masking, crosstalk can be reduced by suppressing hot pixels, which increase DCR in adjacent pixels. However, pixel masking is not available in SS3. Hence, crosstalk could only be mitigated here using computational estimation techniques. To compensate for these effects, 255 or 1024 one-bit images were captured with a total gate exposure of 20 ms. In post-processing, raw images are first corrected for pile-up, followed by interpolation of hot pixels and DCR subtraction. Additional post-processing steps can be applied, such as subtracting the median DCR from each pixel to distinguish crosstalk from normal dark counts and omitting pixels adjacent to hot pixels [1].

**Temporal Gating** is a mechanism in which a SPAD records a photon only if it arrives within a predetermined time window. This window is controlled using a pulse generated by an external source, typically referenced to the laser pulse. However, the rise and fall edges of the gate introduce artifacts in the raw data, posing limitations on the overall system performance. Gate skew, which refers to the time delay between gate edges across the array, and gate edge jitter, the temporal uncertainty of the gate edge position for a single pixel, cannot be corrected due to their stochastic nature and impact the signal-to-noise ratio (SNR).

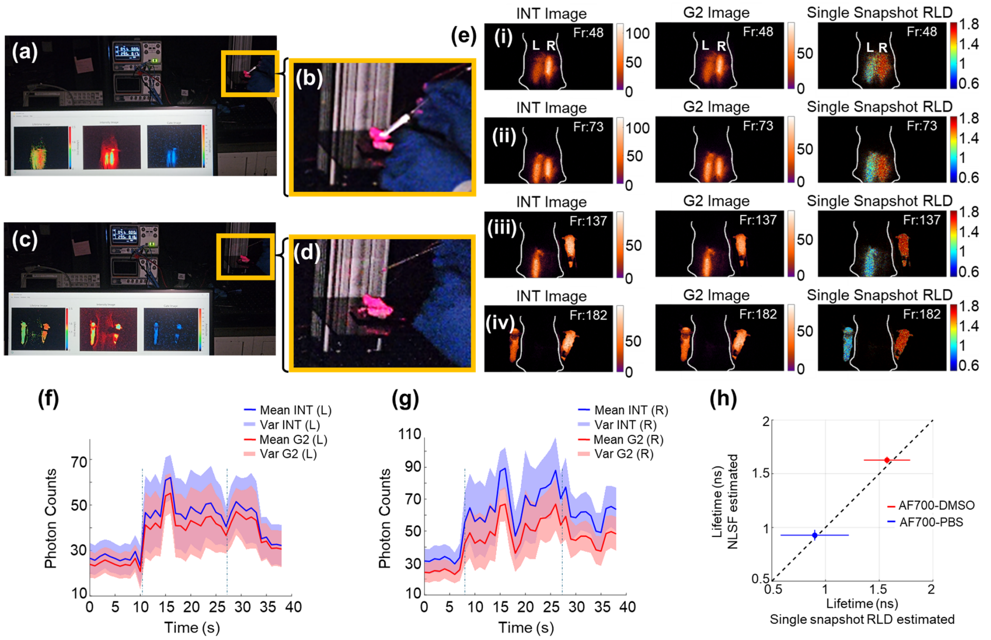


Fig. 4**: Mouse phantom mock surgery.** (a) and (c) show real-time procedure snapshots, with two representative frames selected from the entire procedure. (b) and (d) display enlarged regions corresponding to (a) and (c), respectively. (e)(i–iv) illustrate the step-by-step process of removing the upper layer, exposing the embedded tubes, and extracting them. From left to right, single snapshot images captured in the INT and G2 channels are presented, with fluorescence lifetime computed using these channel data. (f) and (g) show the temporal variation of the INT and G2 signals throughout the procedure, respectively. (h) presents the fluorescence lifetime estimation of the embedded fluorophores, AF700-PBS (left) and AF700-DMSO (right), using a full decay fitting approach (NSLF).

### II. Acquisition and Analysis Software

For real-time acquisition and RLD processing, we modified SwissSPAD Live [5] software, an open-source LabVIEW-based program for data collection and analysis with SwissSPAD2 (SS2) and SS3 time-gated SPAD cameras. It interfaces directly with FPGA firmware (bitfiles), providing a flexible configuration of gate parameters, exposure sequences, and gating protocols, while an auto-reset mechanism maintains continuous operation if an FPGA timeout occurs. The graphical user interface includes controls for microlens usage, reversed gate shifting, and 10-bit dynamic range, as well as settings for gate offset, gate width, gate shift, and buffer size. Laser periods can be defined to align gating with the excitation source (pulsed laser in this case), and a real-time preview feature in software aids in alignment and optical focus on the sample field-of-view. The software also employs hot pixel removal to improve image clarity. Data transfer from the FPGA occurs asynchronously, with visual cues to track progress and detect potential issues.

In this study, SwissSPAD Live was customized to integrate the RLD algorithm for fluorescence lifetime estimation, triggered automatically after the designated gate is captured. A MATLAB script node was used to run the function containing the RLD algorithm, without utilizing any of LabVIEW's parallel processing or LUT (Look-Up Table) optimization features. This limits the potential for increasing the frame rate. The SS3 camera's dual-gate feature captures both a selected gate and its intensity, and the gate of interest can be changed via the settings window. For the RLD experiments, we utilized a gate integration time of 20.451 ms with a gate width of 3 ns. During acquisition, the interface displays second-per-frame information and simultaneously shows the gate image (G2) intensity image (INT), and computed single snapshot lifetime image, allowing immediate assessment and adjustment of experimental conditions.

### III. Non-Linear Least Square Fit (NLSF)

#### A. Iterative Curve Fitting Algorithm

The iterative reconvolution-based curve fitting approach also known as non-linear least square fitting (NLSF) Equation 1 was used for generating the baseline fluorescence lifetime estimation to compare the rapid lifetime determination (RLD) algorithm estimated values for experimental data. For that, the experimental time-resolved data (TPSFs) and their respective instrument response functions (IRFs) were collected from the SwissSPAD3 (SS3) detector. All full laser period TPSFs were fitted using the Levenberg- Marquardt NLSF algorithm implemented in AlliGator software [6]. This method uses periodic or cyclic convolution hence full laser period IRF data was required. Hence, the full-period IRF was used for cyclic convolution with a single bi-exponential periodic decay model. The weighted fit was performed using the minimization function in Equation 2.

$$\begin{cases} \S_T(t) = B + I(t)_T|_{t_0} \circledast F_T(t) \\ F_T(t) = I_0(t) * A_0 e^{-\frac{t}{\tau}} \end{cases} \tag{1}$$

Here, $T$ represents the laser period; $F_T$ (t) periodic sample decay, $t_0$ temporal offset parameter for IRF, $\circledast$ represents cyclic convolution; $A_0$ and $\tau$ are the amplitude and lifetime of fluorophore. B is a baseline parameter accounting for residual uncorrelated background. $\S_T(t)$ is the computed T-periodic fluorescence decay.

$$\chi^2 = \frac{1}{\text{dof}} \sum_{p=1}^{G} \frac{(F_T(t_p) - G_p)^2}{|G_p|} \tag{2}$$

Where dof is the number of independent parameters of the fit, $t_p$ is the $p$th gate location in the laser period, $G$ represents the number of gates, and $G_p$ is the $p^{th}$ gate value. If $G_p = 0$, a weight of 1 replaces the factor $|G_p|$ in Equation 2.

This NLSF process is an iterative fitting algorithm, that optimizes the fitting parameters, depending upon the guess parameters provided by the user. Hence, for large TPSF datasets, the NLSF process can take a relatively long time to extract decay parameters.

### IV. Rapid Lifetime Determination

#### A. General Functions

Dirac delta function ($\delta$ distribution)

$$\delta(x) = \begin{cases} 0, & x \neq 0, \\ \infty, & x = 0, \end{cases} \tag{3}$$

such that

$$\int_{-\infty}^{+\infty} \delta(x)\, \mathrm{d}x = 1. \tag{4}$$

#### B. Relation between Convolution ($*$) and Cyclic Convolution ($\circledast$)

Let $f(t)$ and $g(t)$ be non-periodic functions. The $T$-periodic summation of $f(t)$ and $g(t)$ is defined as the infinite sum of shifted versions of the original functions:

$$f_T(t) = \sum_{i=-\infty}^{+\infty} f(t - iT), \quad g_T(t) = \sum_{i=-\infty}^{+\infty} g(t - iT). \tag{5}$$

The convolution of $f$ and $g_T$ is expressed as [7]:

$$\begin{aligned} f * g_T(t) &= \int_{-\infty}^{+\infty} f(u) g_T(t-u)\, du \\ &= \sum_{i=-\infty}^{+\infty} \int_{-\infty}^{+\infty} f(u) g_T(t - u + iT)\, du \\ &= \sum_{i=-\infty}^{+\infty} \int_{0}^{T} f(v + iT) g_T(t - v)\, dv \\ &= \int_0^T f_T(v) g_T(t-v)\, dv = f_T \circledast g_T(t). \end{aligned} \tag{6}$$

This establishes the identity between the convolution product involving an integral with infinite bounds of a non-periodic function with a $T$-periodic one and the cyclic (or circular) convolution product, denoted by a $\circledast$ symbol, involving an integral of two $T$-periodic functions over a single period $T$.

Note that the convolution product as defined in the last line of Eq. (1) is commutative:

$$\begin{aligned} f_T \circledast g_T(t) &= g_T \circledast f_T(t) \\ &= f * g_T(t) \\ &= f_T * g(t) \\ &= g_T * f(t) \\ &= g * f_T(t). \end{aligned} \tag{7}$$

#### C. Look-up table (LUT), Gate Selection and Offset correction

For fast lifetime computation, a look-up table (LUT) was created by mapping lifetime decay values ($\tau$) to the G2 to INT signal ratio. The following two methods can be used for this.

**Analytical Method** – This method is discussed in **Section 4.3.1** of the Main Document and assumes a square gate and the knowledge of the gate offset $s$ with respect to the laser pulse (which can be estimated experimentally). The relationship between the fluorescence lifetime $\tau$ and the ratio of the INT channel to the G2 channel signal is defined using **Equation 18**. Because **Equation 18** for $\tau$ cannot be inverted algebraically, an iterative numerical approach is needed to extract $\tau$ from the measured INT and G2 channel values. Rather than performing this calculation each and every time, the ratio **Equation 18** is calculated for a range of $\tau$ values relevant for the samples at hand and used as a lookup table (LUT) to which to compare any measured ratio, and obtain the corresponding lifetime value.

**Interpolation Method** – In the case where the gate cannot be assumed to be square-shaped, an interpolation method based on a set of numerically estimated G2 and INT values (or G2/INT ratios) calculated for representative lifetimes $\tau$ using the experimentally measured IRF can be used instead.

In this method, a mono-exponential decay of a pure sample with lifetime decay values ranging from 300 ps to 3000 ps was simulated and convolved with the experimental instrument response function (IRF) obtained at 700 nm and 750 nm, according to **Equation 10** of the Main Document (see Figure 6 and Figure 7 (c) and (d)). The ratio of G2 and INT signals for each gate of the temporal decay, across all lifetime ranges, was used to generate the LUT, as shown in Figure 6 (a) and Figure 7 (a).

The selection of the optimal gate width and location within the laser period for single-snapshot data acquisition depends on the range of lifetimes of interest.

The principle is to obtain the highest variation in the ratio **Equation 18** across the full lifetime range. For instance, in the RPI measurements, based on simulated data using the experimental IRF, the preferred gate at 700 nm was gate number 40 (Figure 6(b)), while at 750 nm, it was gate number 50. However, both gates fall near the beginning of the rising edge or the end of the falling edge, where photon counts are very low. In raw data, noise dominates at such low photon counts. Therefore, a gate with significantly higher photon counts and sufficient ratio variation was chosen. Ultimately,

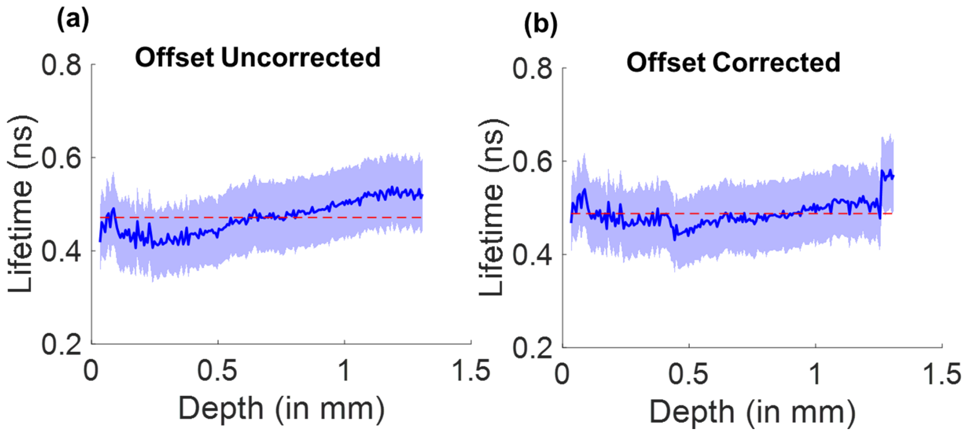


Fig. 5: $45^0$ **mesoscopic lightsheet RLD correction with depth (tumor spheroid Figure. 5(f) main manuscript** (a) The offset uncorrected lifetime estimation with depth. (b) The offset corrected lifetime estimation with depth.

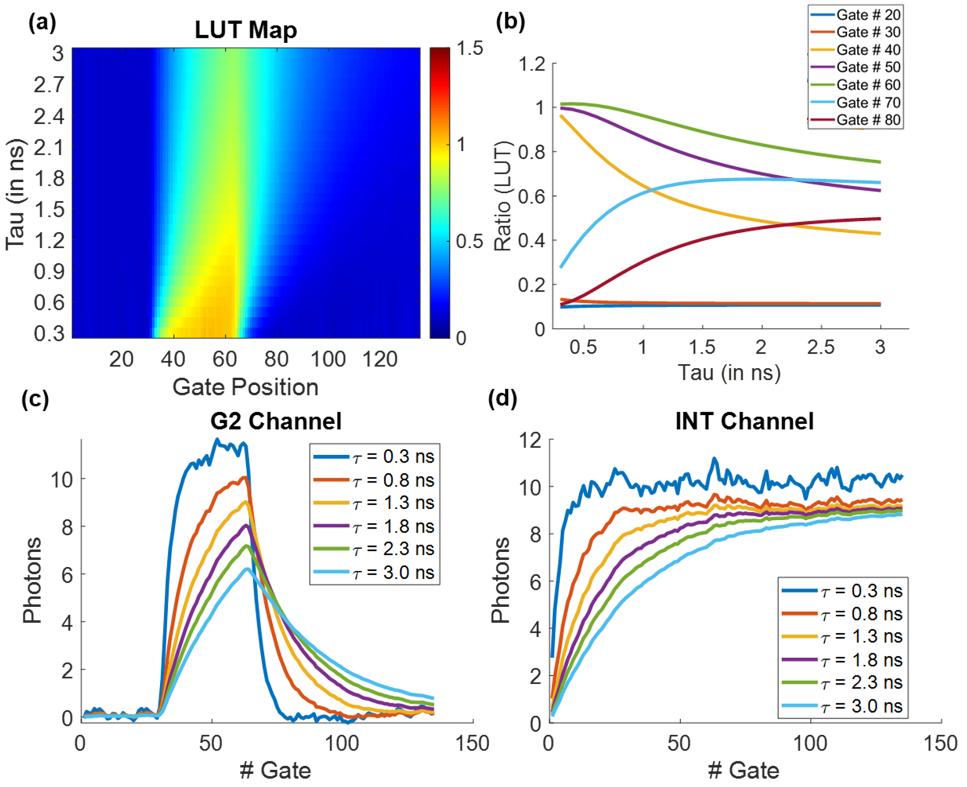


Fig. 6: **Look-up table and Gate selection (analytical Method) for randomly selected pixel at 700 nm** (a) the look-up table for lifetime decay rates and gate position, (b) optimized gate selection for working on various lifetime ranges (c) variation of time-resolved decay using fixed boxcar gate (G2 Channel) with different lifetime ranges (d) variation of time-resolved decay using full aperture (INT Channel) with different lifetime ranges

for 700 nm and 750 nm, gates between 50–60 and 30–40, respectively, were selected.

### V. Deep Learning-Enhanced Single-Snapshot FLI

Single-snapshot-based RLD method offers significant computational benefit but are prone to estimation variation in low photon count regimes due to dominating uncorrectable noise from electronics (see subsection I-B). To overcome this challenge, we developed a deep learning model based on widely used U-Net architecture [8] which is well-suited for image-to-image learning tasks and capable of minimizing RLD estimation variation and enhancing precision. Our DL model learns the noise characteristics introduced by aforementioned artifacts and efficiently corrects the estimation precision. The model is designed to take three feature maps as input: the INT channel image, the G2 channel image, and the single-snapshot computed FLI map as a 3D tensor $X$ of shape $(H, W, C_{in})$ where $H$ and $W$ are the height and width of feature maps,

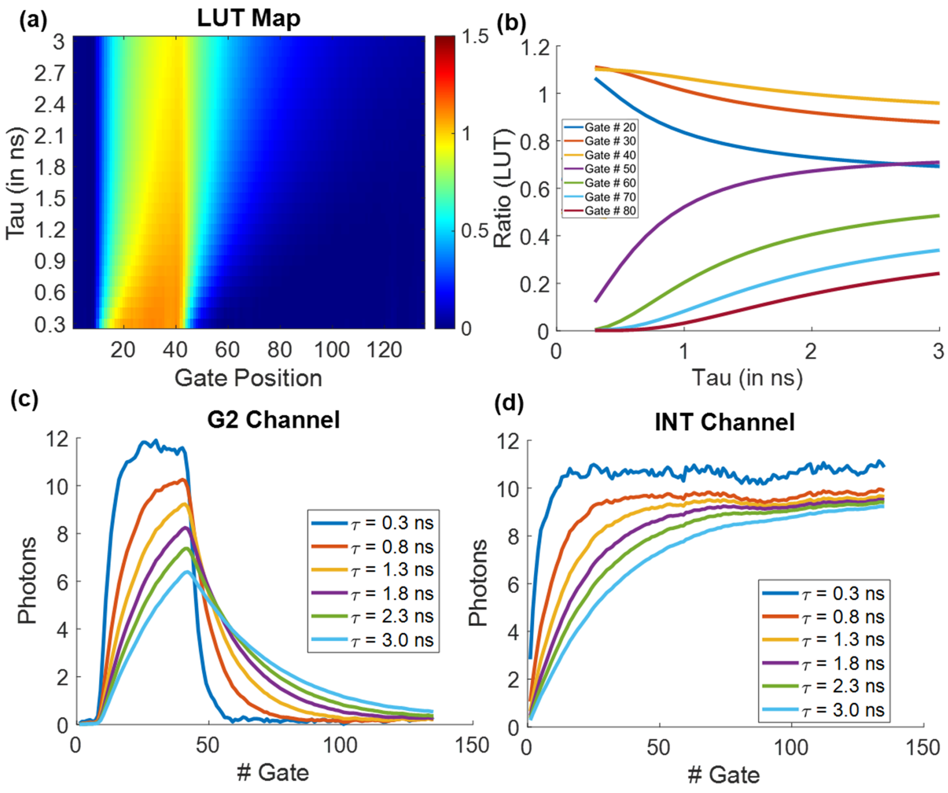


Fig. 7: **Look-up table and Gate selection (Analytical Method) for randomly selected pixel at 750 nm** (a) the look-up table for lifetime decay rates and gate position, (b) optimized gate selection for working on various lifetime ranges (c) variation of time-resolved decay using fixed boxcar gate (G2 Channel) with different lifetime ranges (d) variation of time-resolved decay using full aperture (INT Channel) with different lifetime ranges

and $c_{in}$ is the channels in this case $C_{in}$ is 3.

#### A. U-Net Building Blocks

The following components are used as building blocks for our model:

**Conv2D**$(X, W)$ denotes the 2D convolution operation. For an input tensor $X$ of shape $(H, W, C_{in})$ and a kernel tensor $W$ of shape $(k_H, k_W, C_{in}, C_{out})$, the output $Y$ of the convolution is given by:

$$Y_{h,w,c_{\text{out}}} = \sum_{c_{\text{in}}=1}^{C_{\text{in}}} \sum_{i=1}^{k_H} \sum_{j=1}^{k_W} X_{h+i-1,\, w+j-1,\, c_{\text{in}}} \cdot W_{i,j,c_{\text{in}},c_{\text{out}}} \tag{8}$$

where $h$ and $w$ are the output height and width indices, and $c_{out}$ is the output channel index. Padding and stride are assumed to be handled within this notation.

**Upsampling2D (Nearest-Neighbor)** Let's assume an input tensor X with dimensions (H, W, C) and let's say the up-sampling factor is $(s_h, s_w)$. The output tensor $Y$ will have dimensions $(s_h H, s_w W, C)$. For nearest-neighbor upsampling, the output value $Y(i, j, c)$ is determined by:

$$Y(i,j,c) = X\Big(\left\lfloor \tfrac{i}{s_h} \right\rfloor,\ \left\lfloor \tfrac{j}{s_w} \right\rfloor,\ c\Big). \tag{9}$$

where:

- $i$ and $j$ are the coordinates of the output pixel.
- $c$ is the channel index.
- $\lfloor \cdot \rfloor$ denotes the floor function (rounding down to the nearest integer).

**GroupNorm**$(X, G)$ denotes the group normalization operation. For an input tensor $X$ of shape $(H, W, C)$, the group normalization is given by:

1) Reshape the image $X$ to $(G, \frac{C}{G}, H, W)$.
2) Compute the mean $\mu_{g,h,w}$ and standard deviation $\sigma_{g,h,w}$ for each group $g$ and spatial location $(h, w)$:

$$\mu_{g,h,w} = \frac{1}{\frac{C}{G}} \sum_{c'=1}^{\frac{C}{G}} X_{g,c',h,w} \tag{10}$$

$$\sigma_{g,h,w} = \sqrt{\frac{1}{\frac{C}{G}} \sum_{c'=1}^{\frac{C}{G}} (X_{g,c',h,w} - \mu_{g,h,w})^2 + \epsilon}, \tag{11}$$

where $\epsilon$ is a small constant for numerical stability.

3) Standardize the image:

$$\hat{X}_{g,c',h,w} = \frac{X_{g,c',h,w} - \mu_{g,h,w}}{\sigma_{g,h,w}}. \tag{12}$$

4) Scale and shift the standardized input using learnable parameters $\gamma$ and $\beta$:

$$\text{GroupNorm}(X)_{g,c',h,w} = \gamma_{c'}\, \hat{X}_{g,c',h,w} + \beta_{c'}. \tag{13}$$

5) Reshape the image back to $(H, W, C)$.

**Dropout**$(X, d)$ denotes the dropout operation, where $d$ is the dropout rate. For each element $X_{b,h,w,c}$ in the input tensor $X$:

$$\text{Dropout}(X)_{b,h,w,c} = \begin{cases} 0, & \text{w.p. } d, \\ \dfrac{X_{b,h,w,c}}{1-d}, & \text{w.p. } 1-d. \end{cases} \tag{14}$$

**Activation Function** For this application we used *Mish* activation function [9] represented as $f(.)$ defined as follows:

$$f(x) = x \tanh(\zeta(x)) \tag{15}$$

$$\zeta(x) = \ln(1 + e^x), \tag{16}$$

where $\zeta(x)$ is the softplus activation.

**Residual Block (ResBlock)** This block processes an input tensor $X$ of shape $(H, W, C_{in})$ where $H$ and $W$ are the height and width of the map, respectively, and $C_{in}$ is the number of input channels. $R$ is the residual tensor defined as

$$R = \begin{cases} X, & \text{if } C_{in} = 3 \\ \text{Conv2D}(X, W_1), & \text{if } C_{in} \neq 3 \end{cases}, \tag{17}$$

where $W_1$ is a kernel of size (1,1) and $d$ is set to 0.05. $Y_{out}$ is the output of the residual block and is computed as follows

$$Z = \text{Conv2D}\left(f\left(\text{Dropout}\left(\text{GroupNorm}(X), d\right)\right), W_3\right) \tag{18}$$

$$\tilde{Z} = \text{Dropout}\left(\text{GroupNorm}(Z), d\right) \tag{19}$$

$$Y = \text{Conv2D}(f(\tilde{Z}), W_3) \tag{20}$$

$$Y_{\text{out}} = Y + R \tag{21}$$

The ResBlocks help stabilize the training of deep networks by allowing gradients to flow more easily through the network [10].

We implemented a U-Net architecture, characterized by its encoder-decoder structure. The encoder progressively extracts features using "Down Blocks" and refines them through residual "Mid Blocks". The decoder then upsamples these features via "Up Blocks", with a final (1,1) Conv2D layer to generate the output with the desired number of channels. The following sections provide a detailed explanation of the encoder and decoder components.

#### B. Model Architecture

**Down Block** The contracting path repeatedly applies convolutions followed by non-linear activation. This is often done with two consecutive convolutions.

$$\begin{aligned} \tilde{Z}_{down-block} &= f(\text{Conv2D}(X, W_3)) \\ Z_{down-block} &= f(\text{Conv2D}(\tilde{Z}_{down-block}, W_3)) \end{aligned} \tag{22}$$

GroupNorm followed by Dropout were used after Conv2D but before *mish* $f(.)$ activation function. GroupNorm is used to stabilize the training process as it normalizes the features within channels, unlike batch normalization which use normalization across the entire batch. The computation of GroupNorm remains independent of batch sizes. The contracting path repeatedly applies these operations, reducing the spatial dimensions of the feature maps.

**Mid Block** The Mid Block extract the feature information between channels. This section uses repeated ResBlocks. In this DL model two ResBlocks were used for this operation for $Z_{mid-block}$.

$$\begin{aligned} \tilde{Z}_{mid-block} &= \text{ResBlock}(Z_{down-block}) \\ Z_{mid-block} &= \text{ResBlock}(\tilde{Z}_{down-block}) \end{aligned} \tag{23}$$

**Up Block** The expansive path consists of upsampling followed by concatenation with the corresponding feature map from the contracting path and convolutions.

The operations can be described as follows:

1) **ResBlock and Concatenation:**

$$\tilde{Z}_{\text{up-block}} = \text{ResBlock}\big(\text{Concat}(Z_{\text{mid-block}})\big) \tag{24}$$

This concatenates the upsampled feature map with the corresponding feature map from the encoder, providing high-resolution features.

2) **Upsampling:**

$$Z_{\text{up-block}} = f\Big(\text{Conv2D}\big(\text{Upsampling2D}(\tilde{Z}_{\text{up-block}})\big), W_3\Big) \tag{25}$$

This refines the feature maps.

The expansive path repeatedly applies above two operations, increasing the spatial dimensions and decreasing the number of feature channels.

$$Y_{output} = \text{Conv2D}\big(f\big(\text{GroupNorm}(Z_{up-block}), d\big), W_3\big) \tag{26}$$

The final layer uses kernel size of a $(3 \times 3)$ to map the desired number of output classes.

The ResBlocks are implemented at each resolution level to improve the model's ability to learn both fine-grained and large-scale features. Each ResBlocks consists of two convolutional layers with group normalization and dropout applied in between. Group normalization was chosen over batch normalization to accommodate smaller batch sizes more flexibly [11], and dropout was used to mitigate overfitting.

In the encoder, at each downsampling step, the spatial resolution is halved while the number of feature channels is increased to capture higher-level abstractions. The decoder upsamples the feature maps and concatenates them with the corresponding encoder outputs, allowing the model to fuse fine-scale details with deeper, context-rich representations [8]. Each decoder stage applies the same residual processing as in the encoder, ensuring consistent feature extraction across scales. Finally, the model outputs a single-channel map at the original resolution, which represents the enhanced fluorescence lifetime map. By integrating elements from established U-Net variants [12] and adopting group normalization for stability under varying batch sizes, our architecture balances the preservation of local details with robust feature abstraction.

The model was trained using the experimental data captured in multiple settings camera settings. we acquired both single-shot data acquisition, INT image and G2 image and RLD computed FLI and full time-gated measurements to generate NLSF-based lifetime map. For the lack of absolute reference

NLSF method was considered as ground truth. Data were collected from both macroscopic and mesoscopic imaging setups as described earlier, and additional data augmentation method was applied to further expand the training set. The synthetics data was also simulated as per previously published work ( [13]). The model was trained with 1000 sets of images. Model training was performed on a Windows computer equipped with an NVIDIA RTX 3090, using early stopping to prevent overfitting, with 20% of the data allocated for testing and another 20% for validation. The inference time for image size $250 \times 484$ was $\sim 400$ ms.
